# Context-Dependent Enhancer Function Revealed by Targeted Inter-TAD Relocation

**DOI:** 10.1101/2022.01.19.476888

**Authors:** Christopher Chase Bolt, Lucille Lopez-Delisle, Aurélie Hintermann, Bénédicte Mascrez, Antonella Rauseo, Guillaume Andrey, Denis Duboule

## Abstract

The expression of genes with a key function during development is frequently controlled by large regulatory landscapes containing multiple enhancer elements. These landscapes often match Topologically Associating Domains (TADs) and sometimes integrate range of similar enhancers, thus leading to TADs having a global regulatory specificity. To assess the relative functional importance of enhancer sequences *versus* the regulatory domain they are included in, we set out to transfer one particular enhancer sequence from its native domain into a TAD with a closely related, yet different functional specificity. We used *Hoxd* genes and their biphasic regulation during limb development as a paradigm, since they are first activated in proximal limb cells by enhancers located in one TAD, which is then silenced at the time when the neighboring TAD starts to activate its enhancers in distal limb cells. We introduced a strong distal limb enhancer into the ‘proximal limb TAD’ and found that its new context strongly suppresses its distal specificity, even though it continues to be bound by HOX13 transcription factors, which normally are responsible for this activity. Using local genetic alterations and chromatin conformation measurements, we see that the enhancer is capable of interacting with target genes, with a pattern comparable to its adoptive neighborhood of enhancers. Its activity in distal limb cells can be rescued only when a large portion of the surrounding environment is removed. These results indicate that, at least in some cases, the functioning of enhancer elements is subordinated to the local chromatin context, which can exert a dominant control over its activity.

## INTRODUCTION

Genes with important functions during vertebrate development are frequently multifunctional, as illustrated by their pleiotropic loss-of-function phenotypes. This multi-functionality generally results from the accumulation of tissue-specific enhancer sequences (Banerji et al., 1981) around the gene. At highly pleiotropic loci, the collection of enhancers can be distributed across large genomic intervals, which are referred to as regulatory landscapes (Bolt and Duboule, 2020; Spitz et al., 2003). Such landscapes often correspond in linear size to Topologically Associating Domains (TADs)(Dixon et al., 2012; Nora et al., 2012; Sexton et al., 2012), defined as chromatin domains wherein the probability of DNA-to-DNA interactions is higher than with neighboring domains. These collections of enhancers likely reflect an evolutionary process whereby the emergence (or modification) of morphologies was accompanied by novel regulatory sequences with the ability to be bound by tissue specific factors, and hence alter the transcription of nearby target genes (Carroll et al., 2008).

This model, however, does not explicitly account for the potential influence of local genomic context to modulate interactions between promoters and nascent enhancers, thus adding another layer of complexity to our understanding of developmental gene regulation. For instance, many developmental gene loci are initially decorated with the chromatin mark H3K27me3 deposited by the Polycomb complex PRC2, before their activation (Bernstein et al., 2005), possibly as a way to maintain silencing or limit transcription to very low levels. These chromatin modifications disappear along with gene activation and are re-deposited once the gene and its enhancers are progressively decommissioned, thus preventing inappropriate transcription that may cause severe developmental problems (see Schuettengruber et al., 2017). This indicates that enhancer or promoter function can be affected by their local chromatin environments that may amplify both repressive or activating cues.

In the case of large regulatory landscapes containing many enhancers with similar tissue specificity, the question arises as to how their activation and decommissioning are coordinated and enforced over very large genomic intervals to overcome any contradictory inputs. Two scenarios can be considered in this context. The first possibility is that each enhancer sequence within the landscape independently responds to information delivered solely by the factors bound to it in a sequence-specific manner. Such a mechanism would act entirely in *trans* and its coordination would be ensured by the binding of multiple similar positive or negative factors. In the second scenario, a higher level of regulation is imposed by the general environment of the landscape, for example corresponding to a TAD-wide positive or negative regulation, which would tend to dominate individual enhancer-specific controls in order to minimize dangerous misexpression events.

To address this issue, we used the development of the vertebrate limb as a paradigm and, in particular, the essential function of *Hox* genes in patterning and producing the main pieces of tetrapod appendages (Zakany and Duboule, 2007). The emergence of the tetrapod limb structure was a major evolutionary change that facilitated animal migration onto land. A key step in this process was the acquisition of a fully developed distal part (hands and feet), articulating with the more ancestral proximal parts of the limbs (e.g. the arm and the forearm). During development, the distal piece requires the specific function of several key genes, amongst which are *Hoxa13* and *Hoxd13*. While the absence of either gene function induces a moderate phenotype in hands and feet (Dolle et al., 1993; Fromental-Ramain et al., 1996), the double loss-of-function condition leads to the agenesis of these structures (Fromental-Ramain et al., 1996) thus suggesting a critical function for these two genes both during the development of distal limbs and its evolutionary emergence (see Woltering and Duboule, 2010). In this context, the regulatory mechanisms at work to control expression of both genes in most distal limb cells were carefully analyzed either for *Hoxa13* (Berlivet et al., 2013; Gentile et al., 2019), or for *Hoxd13* (Montavon et al., 2011; Spitz et al., 2003), and found to be quite distinct from one another.

In the case of *Hoxd* genes, the evolutionary co-option of *Hoxd13* into both the distal limbs and the external genitals was accompanied by the emergence of a novel TAD (C-DOM) containing multiple enhancers specific to either tissue or common to both (Acemel et al., 2016; Amândio et al., 2020; Montavon et al., 2011; Woltering et al., 2014). This TAD is inactive until late in development, when limb cells with a distal fate start to appear. In contrast, the expression of other *Hoxd* genes (*Hoxd9*, *Hoxd10* and *Hoxd11*), which are all critical for the formation of more proximal parts of the limb (Davis et al., 1995), are controlled by enhancers acting earlier and located in the adjacent TAD (T-DOM)(Andrey et al., 2013). The *HoxD* cluster thus lies in between these two TADs and contains several CTCF binding sites that create a boundary between the two adjacent chromatin domains, insulating *Hoxd13* and *Hoxd12* from the rest of the gene cluster (Rodriguez-Carballo et al., 2017). In the early stages of limb bud formation, *Hoxd9*, *Hoxd10,* and *Hoxd11* are activated by the regulatory landscape located telomeric to the cluster (T-DOM) leading to the patterning and growth of long bones of the arms and legs. Subsequently, in a small portion of cells at the posterior and distal end of the early limb bud, enhancers located in the centromeric regulatory landscape (C-DOM) are switched on, driving expression of 5’-located *Hoxd* genes in the nascent hands and feet (Andrey et al., 2013).

The operations of these two TADs are mutually exclusive and as soon as C-DOM starts to upregulate expression of *Hoxd13* in distal cells, enhancers within T-DOM are decommissioned and a large part of this chromatin domain becomes decorated by H3K27me3 marks (Andrey et al., 2013). This switch in TAD implementation involves the HOX13 proteins themselves since their production in response to C-DOM enhancers participate in the repression of T-DOM enhancers, likely through direct binding (Beccari et al., 2016; Sheth et al., 2016). In parallel, HOX13 proteins positively regulate C-DOM enhancers, thus re-enforcing the functional switch between the two TADs. In the absence of both HOXA13 and HOXD13, C-DOM is never activated whereas T-DOM enhancers continue to function due to the absence of both decommissioning and H3K27me3 coverage (Beccari et al., 2016). Therefore, in this context, HOXD13 shows properties of both a transcriptional activator and a transcriptional repressor at different times and in different chromatin environments.

At the time when *Hoxd13* becomes activated by distal limb (digit) enhancers, it is critical that all proximal limb (forearm) enhancers are rapidly switched off, to prevent that the latter may act on the former, a situation that was shown to be detrimental to limb morphology (Bolt et al., 2021). This bimodal regulatory situation thus provides a paradigm to assay for the presence of a ‘context-dependent’ repression of enhancer activity. Accordingly, we set out to introduce into the proximal limb-specific T-DOM domain, the strongest distal enhancer normally working within C-DOM to activate *Hoxd13* in digit cells, asking whether this distal limb regulatory sequence would still be able to exert its function when relocated into a TAD where limb enhancers are being repressed in distal cells. We report that this enhancer element, which is functionally very penetrant when introduced at various random positions by non-targeted transgenesis, loses most of its distal limb activity when recombined within T-DOM, even though it continues to recruit the HOX13 factors, which are essential for its function in distal limb cells. We further show that part of the distal limb activity is restored when large portions of T-DOM are deleted *in-cis*. We conclude that the function of this enhancer is inhibited by an *in cis* mechanism acting at the level of an entire chromatin domain, suggesting the existence, in this particular case, of a level of regulation higher than that of the enhancer sequences themselves.

## RESULTS

### A distal-limb specific enhancer

To evaluate how a tissue specific enhancer would behave when relocated into a different chromatin and regulatory context, we searched for a candidate enhancer element that would display the strongest possible specificity for developing distal limb cells at the *HoxD* locus. We performed ATAC-seq in wild type E12.5 distal limb cells and compared the signals with a previously reported H3K27ac ChIP-Seq dataset (Rodriguez-Carballo et al., 2017). Because distal limb enhancers located at *Hox* loci were shown to colocalize with the binding of HOX13 proteins as assayed in ChIP-seq experiments (Beccari et al., 2016; Sheth et al., 2016), we used a CUT&RUN approach for both HOXA13 and HOXD13 (together referred to as HOX13) to more precisely delineate enhancer elements controlled, at least in part, by these transcription factors.

We identified a small element within a region previously described as Island II, one of the islands of the C-DOM regulatory archipelago (Lonfat et al., 2014; Montavon et al., 2011). This element was strongly bound by both transcription factors and accessible as judged by ATAC-Seq, suggesting that it is an active enhancer element in distal limb cells and that it controls the expression of the most posterior *Hoxd* genes there (Figure 1A). Upon closer inspection, we found that the peak from CUT&RUN experiments matched precisely with the peak observed in a previously reported E11.5 whole limb bud ChIP-Seq experiment for HOXA13 and HOXD13 (Figure 1B, dark grey lines)(Sheth et al., 2016). It also matched a small DNA fragment that is highly conserved across tetrapods but thus far absent from all evaluated fish genomes including that of coelacanth (Figure S1A; dashed box). This element had all of the hallmarks of an active distal limb enhancer controlled by HOX13 transcription factors and hence we called it enhancer element II1.

**Figure 1.**
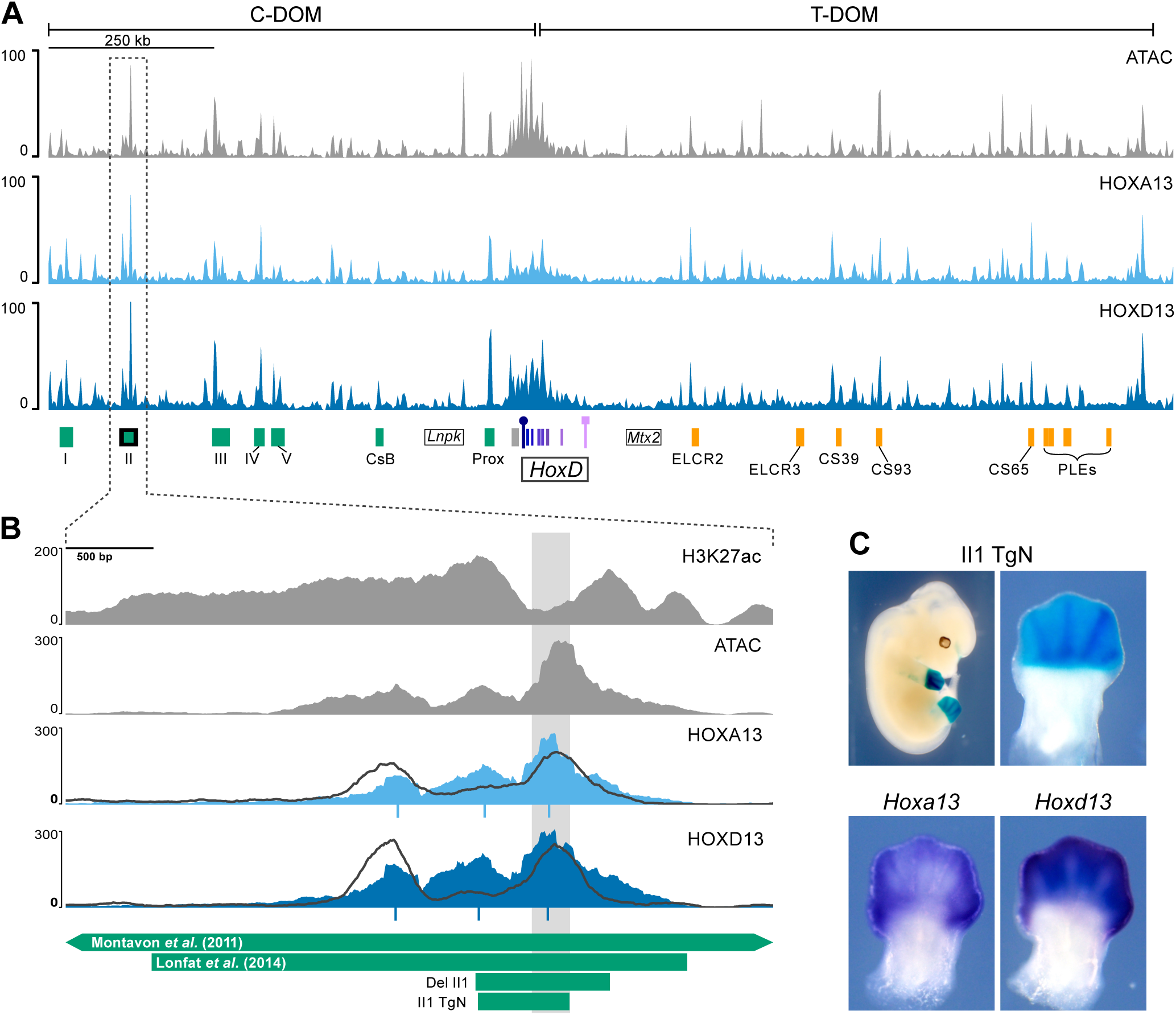
Identification of a distal limb bud specific enhancer. **A**. ATAC-seq profile (top) and binding profiles of both the HOXA13 (middle) and HOXD13 (bottom) transcription factors by CUT&RUN, using E12.5 wildtype distal forelimb cells and covering the entire *HoxD* locus, including the two flanking TADs C-DOM and T-DOM (mm10 chr2:73950000-75655000). Green rectangles below are distal limb enhancers in C-DOM and orange rectangles are proximal limb enhancers in T-DOM. The *Hoxd* gene cluster is indicated with a box at the center and the *Lnpk* and *Mtx2* genes are indicated as rectangles with black borders. The *Hoxd13* gene is on the centromeric end of the cluster and indicated by a purple box with a circle on the top, whereas *Hoxd1* is telomeric and indicated by a square on top. The C-DOM Island II enhancer (Montavon et al., 2011) is a green rectangle with a black border and its corresponding signals are indicated by a dashed vertical rectangle. **B**. Magnification of the same tracks as above centered around the Island II enhancer with, in addition, the H3K27ac ChIP-seq signal (top) in E12.5 distal forelimbs (Rodriguez-Carballo et al., 2017). The profiles indicated by the dark grey lines are from ChIP-seq using E11.5 whole limb buds (Sheth et al., 2016) for comparison and are shown for comparison. Below the CUT&RUN profiles are the MACS2 peak summits for the corresponding CUT&RUN samples. The green rectangles below indicate the regions described as Island II in (Montavon et al., 2011) or in (Lonfat et al., 2014). The Del II1 shows the region deleted in this work (Figure S1) and the II1 TgN is the transgene used panel in C. **C.** *LacZ* staining pattern produced by the II1:*HBB:LacZ* (II1 TgN) enhancer reporter transgene at E12.5, showing high specificity for distal limb cells (top). Below are whole-mount *in situ* hybridizations for *Hoxa13* and *Hoxd13* in wild type E12. 5 forelimbs for comparison.

To test if this short element indeed carries the enhancer activity reported for the large versions of Island II (Figures 1B and S1A; green rectangles below), we cloned the sequence (532bp, mm10 chr2:74075311-74075843) and constructed a reporter transgene where the enhancer is located 5’ to the *HBB* promoter and the *LacZ* gene (Figures 1B and S1A; II1 TgN), and injected it into mouse pronuclei. Five founder animals were identified and then crossed with wild types. Embryos were collected at E12.5 and stained for *LacZ*. Four of the founders transmitted the transgene and produced strong distal limb specific staining (Figure 1C, top). One of the four transmitting founders also had low staining levels in nearly the entire embryo and another founder produced staining in the central nervous system. The four animals transmitting limb staining displayed a pattern closely matching the expression domains of both *Hoxd13* and *Hoxa13* (Figure 1C). The variation observed in staining in other tissues likely resulted from different integration sites.

### Deletion of the II1 enhancer sequence

The C-DOM regulatory landscape contains multiple enhancer elements that are necessary to produce a robust activation of *Hoxd* genes in the distal limbs and genitals (Amândio et al., 2020; Montavon et al., 2011) and our ATAC-Seq and CUT&RUN experiments showed that element II1 is among the most strongly bound and accessible elements throughout the C-DOM (Figure 1A). Because of this, we anticipated that it may make a measurable contribution to the transcription of *Hoxd* genes in the distal limb. To determine what effect this element contributes to the global regulatory activity of C-DOM, we used CRISPR/Cas9 to delete the II1 enhancer element (Figures 1B and S1A; green rectangle Del II1). Several founders were obtained and we produced embryos homozygous for this (*HoxD^DelII1^*) deletion. We collected embryos at E12.5, measured their levels of distal limb *Hoxd* mRNAs by RT-qPCR and looked at the transcript distribution by *in situ* hybridization. Using both approaches, we did not observe any significant change, neither in the level of *Hoxd* gene transcription in the distal limbs of embryos missing the II1 enhancer (Figure S1B, C), nor in the spatial distribution. While this result was somewhat surprising given how strong the signal for H3K27ac, ATAC and HOX13 binding are, a similar lack of effect was observed when single genital enhancers were deleted within this same C-DOM TAD. However, removing several such enhancers had a cumulative effect thus demonstrating the regulatory resilience of this regulatory landscape (Amândio et al., 2020).

### Targeted insertion of a C-DOM enhancer into T-DOM

The T-DOM TAD contains multiple enhancers (Figure 2A, orange rectangles), which activate *Hoxd* genes in proximal limb cells, and the deletion of T-DOM abolishes all *Hoxd* gene transcripts from the proximal pieces of the growing limb buds (Andrey et al., 2013; Bolt et al., 2021). In distal limb cells these elements are no longer at work and the entire T-DOM is switched off at the same time as enhancers within C-DOM are activated, including the II1 element. HOX13 proteins bind throughout the T-DOM and are associated with the decommissioning of the proximal limb enhancers. When HOX13 factors are not present the decommissioning does not occur and the ‘proximal’ enhancers continue operating in distal cells (Beccari et al., 2016). In fact, the T-DOM proximal enhancers CS39 and CS65 also operate in distal cells when introduced randomly into the genome as transgenes (Beccari et al., 2016), suggesting that the T-DOM environment represses the function of such sequences in distal cells, and that this mechanism may involve the binding of HOX13 factors.

**Figure 2.**
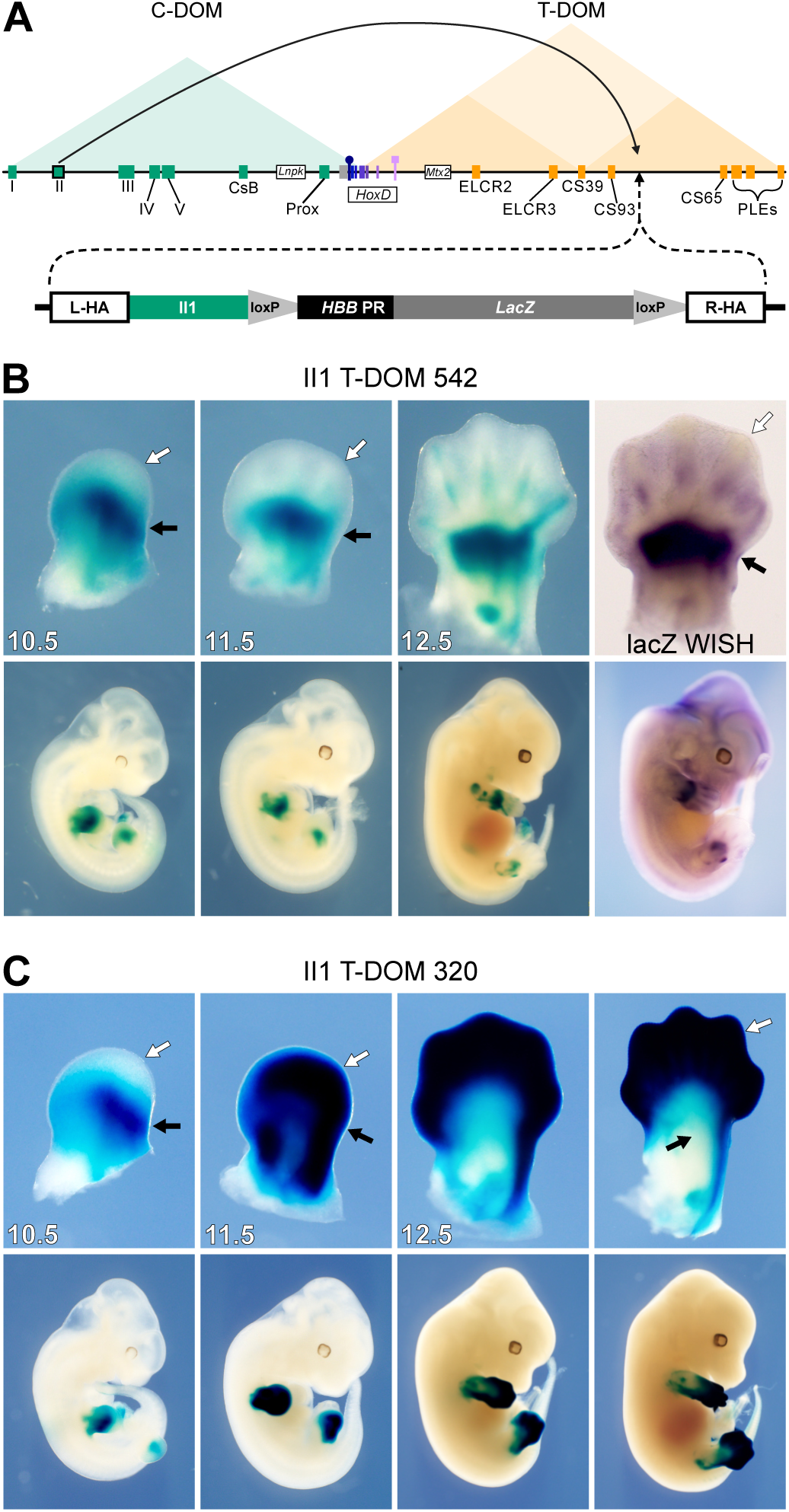
Targeted recombination of the II1 reporter construct into T-DOM. **A**. General scheme of the *HoxD* locus with the two triangles on top showing the extents of both the C-DOM (green) and T-DOM (orange) TADs. Various enhancers are shown either in green (distal limbs) or yellow (proximal limbs) rectangles and the *HoxD* cluster is boxed. The large arrow on top indicates the origin and new location of the island II enhancer transgene into T-DOM. Below is a map of the II1:*HHB:LacZ* construct containing both left (L-HA) and right (R-HA) homology arms, the II1 enhancer element and the *HBB* promoter with a *LacZ* reporter gene. **B**. *B-galactosidase* staining time course and *LacZ* mRNAs (right panel) of the single-copy II1 T-DOM 542 founder line. At E10.5 and 11.5, staining is very strong in the proximal limb (white arrow) while absent in the distal portion (black arrow). By E12.5 weak staining appears in the digit mesenchyme of the distal limb. The WISH for *LacZ* mRNA confirms that the distal limb staining comes from transcription in distal limb cells rather than from stable *B-galactosidase* activity. **C**. *B-galactosidase* staining of the multi-copy II1 T-DOM 320 founder line. Staining is much stronger in this allele and similar to line 542 in E10.5 limb buds. However, by E11.5, strong staining is gained in distal limb cells whereas it disappears from the proximal domain.

To further challenge this repressive effect and see whether it would dominate over the strong distal specificity of an enhancer that normally operates in distal cells, we set up to relocate a single copy of the enhancer II1 into T-DOM. We used the same transgenic construct (Figure 1, II1 TgN) used to test the enhancer element by random insertion transgenesis (Figure 1C), but we attached homology arms to the 5’ and 3’ ends to target insertion to a region of T-DOM in between - yet at a distance from - two strong CS39 and CS65 proximal limb enhancers (Andrey et al., 2013; Beccari et al., 2016). This region was also selected because it had low levels of the Polycomb Group histone H3 modification H3K27me3, such that H3K27me3 short distance spreading (see Cheutin and Cavalli, 2019) would not directly impact the inserted element. Fertilized eggs were injected with both the targeting construct and various CRISPR components (Figure 2A, Table S1).

We identified two founder lines that carried the construct and produced *LacZ* staining. The *HoxD^II1-T-DOM-542^* founder line (‘allele 542’) showed a strong *LacZ* staining in the proximal limb (Figure 2B, black arrows), with very weak staining appearing in the distal limb from E12.5 onwards (Figure 2B, white arrows), limited to a small region of the forming digits. Since the II1 transgene was located within T-DOM, which hosts multiple proximal limb enhancers, the strong proximal limb activity likely resulted from the transgene behaving as an enhancer sensor in these cells. The second line identified (*HoxD^II1-T-DOM-320^*) produced a strikingly different staining pattern (Figure 2C). At E10.5 both the II1 T-DOM 542 and II1 T-DOM 320 lines had strong staining in the proximal limb, but not in the distal limb (Figure 2B, C; compare white and black arrows). Then, at E11.5, while both lines continued to produce strong proximal limb staining, the II1 320 line also had strong distal limb staining (Figure 2C, white arrows). By E12.5 the staining pattern between the lines had almost completely diverged, with II1 T-DOM 542 showing strong proximal and very weak distal staining. In contrast the II1 T-DOM 320 line produced very strong distal staining but it was nearly absent in the proximal limb (Figure 2B, C).

To try to understand this difference and to confirm that the II1 transgene was inserted at the expected position, we performed long-read sequencing with the Oxford Nanopore MinION in conjunction with the nCATS protocol (Gilpatrick et al., 2020) to enrich for sequencing reads covering the region around the insertion site (Figure S2A). In the 542 sample, we were able to detect a sequencing read that extended the length of the transgene, both homology arms, and several kilobases of the region that flanks the insertion (Figure S2B), thus confirming that the 542 allele was a single copy transgene inserted correctly at the right site. Because this enhancer was inserted into T-DOM (hereafter II1 T-DOM) producing the correct allele, we used it for all subsequent experiments. We were not able to completely sequence the II1 T-DOM 320 allele because it contained multiple tandem copies of the insertion, yet we could reconstruct a putative genome based on unique overlapping reads which indicated that a minimum of four copies of the transgene were inserted as a tandem array with multiple orientations (Figure S2C).

From the analysis of these two alleles, we conclude that when inserted as a single copy at the selected position within T-DOM, the distal limb enhancer activity of II1 was almost entirely repressed and thus this sequence behaved there like any other native proximal limb enhancers located in T-DOM (Andrey et al., 2013; Beccari et al., 2016). Since the heterologous promoter trapped the activity of the surrounding proximal limb enhancers, the final pattern appeared exactly as the reverse pattern for this enhancer when operating from C-DOM. This positive response in proximal limb cells also acted as an internal control; the transgene was capable of transcriptional activity, with a capacity for tissue specific expression. The repression from the chromatin environment in distal cells was completely alleviated when the transgene was present in multiple tandem copies, suggesting a potential micro-structure capable of escaping the negative effect of the local TAD environment. The decreased expression of this allele in proximal cells suggested that the array of transgenes was somehow insulated from receiving the influence of T-DOM proximal enhancers.

### HOX13 transcription factors bind to II1 T-DOM in distal limb cells

Despite the observed repression, when the II1 transgene was inserted into T-DOM as a single copy, we scored a very low level of *LacZ* staining in E12.5 distal limb cells, detected only at a late stage and appearing as defined spots in the digit mesenchyme, i.e., in cells that do not reflect well the wild type expression pattern which is more widespread at this stage (Figure 1C and 2B). Because the native II1 enhancer activity seems to be dependent on HOX13 proteins (Desanlis et al., 2020), we looked for both the accessibility of- and HOX13 binding to the II1 element within the context of the inactive T-DOM in distal limb cells, to see if HOX13 factors could still access and bind the relocated enhancer sequence.

We first performed an ATAC-seq on embryos homozygous for the II1 T-DOM single-copy insertion along with wild type controls. We mapped the reads on the mutant genome and used a high mapping quality score (MAPQ30) to ensure that all reads observed on the coverage tracks mapped uniquely to the native II1 element in C-DOM (II1 C-DOM) and the II1 transgene recombined into T-DOM (II1 T-DOM). In wild type control samples, as expected, no ATAC signal was observed over the native II1 C-DOM, neither in proximal limb bud cells, nor in forebrain cells (Figure 3A left). In contrast, a strong ATAC signal appeared over the II1 element in distal limb cells (Figures 1B, 3A; DFL). In the II1 T-DOM transgene, there was a weak signal observed over the *HBB* promoter element in forebrain and proximal limb cells, the latter signal in proximal forelimb cells (PFL) likely reflecting the *LacZ* staining in response to nearby proximal limb enhancers. In the distal forelimb samples (DFL), there was a further increase in signal over the *HBB* promoter, and the accessible region extended over the II1 enhancer portion of the transgene. This signal over the II1 enhancer was nevertheless weaker than that observed on the II1 element located in its native context (Figure 3A, ATAC DFL, compare left and right).

**Figure 3.**
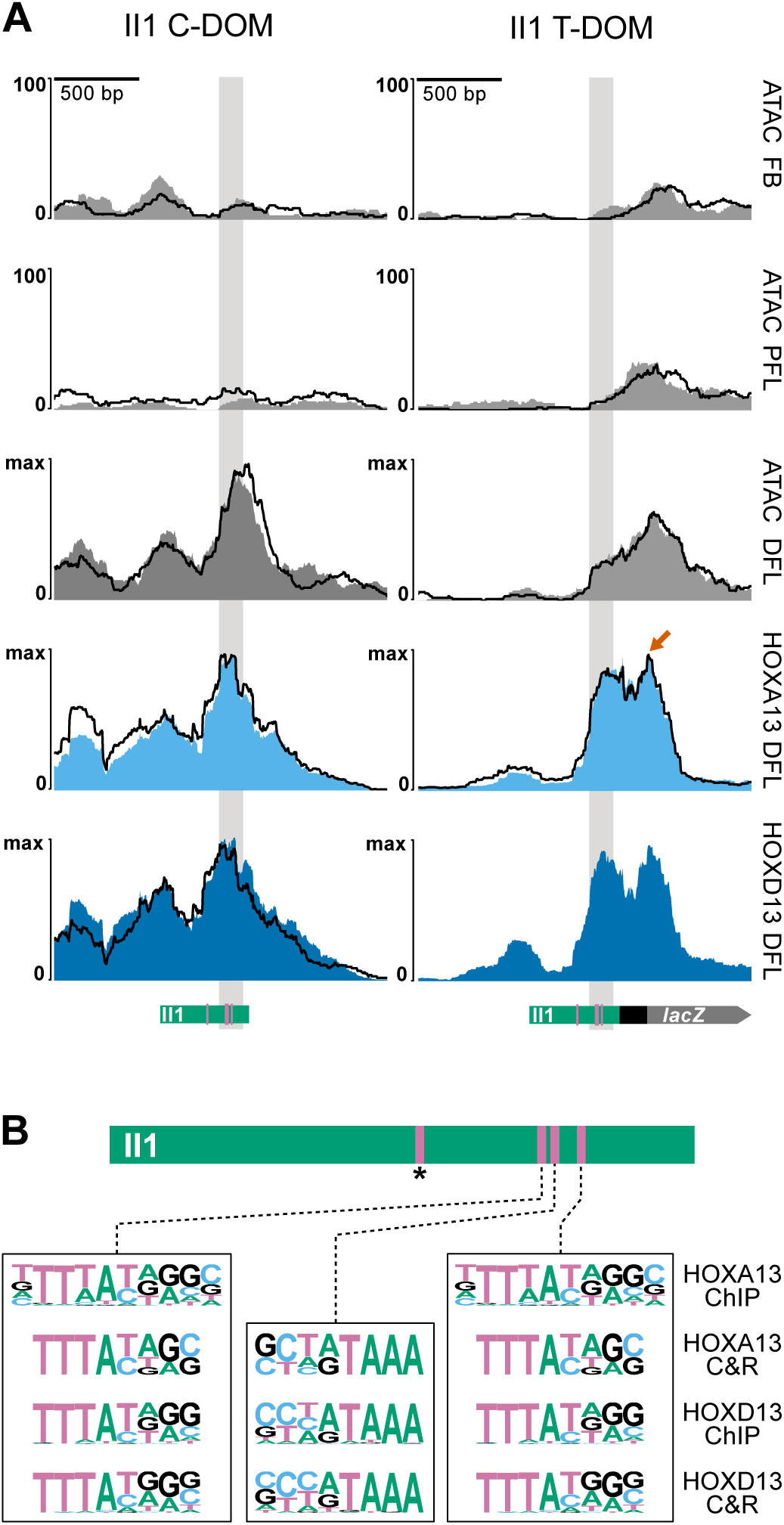
HOX13 proteins bind to the II1 enhancer in T-DOM. **A**. ATAC-Seq and CUT&RUN reads mapped to the II1 enhancer sequence, either in its native environment within C-DOM (left column; mm10 chr2:74074674-74076672), or after its targeted recombination within T-DOM (line 542, right column; mm10 chr2:75268925-75270923). The green rectangles below indicate the extent of the region used for the II1 enhancer element with, in pink, the position of the three HOX13 binding sites. The II1 C-DOM element is not accessible by ATAC-Seq in E12.5 forebrain (FB), nor in proximal forelimb cells (PFL) samples. At E12.5 it becomes highly accessible in distal forelimb cells (DFL) and is strongly bound by HOX13 proteins. The II1 element in T-DOM has low accessibility in the FB and PFL samples, even though there is high transcription of the transgene in PFL. Similar to the II1 element in C-DOM, the II1 enhancer in T-DOM is occupied by HOX13 proteins in distal limb cells. It also shows an additional peak over the *HBB* promoter (orange arrow). This peak is likely a non-specific signal resulting from promiscuous MNase activity used in the CUT&RUN technique. In all samples but HOXD13 in the II1 T-DOM allele, experiments were performed in duplicate. One replicate is plotted as a solid color and the other is shown as a superimposed black line. **B**. On top is a schematic of the II1 enhancer element with the four HOX13 motif indicated as pink bars. The pink bar with an asterisk indicates the motif position that is not in the HOX13 and ATAC peak. At the bottom are the three HOX13 motifs identified by HOMER motif discovery in the CUT&RUN experiments here and the E11.5 whole forelimb ChIP-seq (Sheth et al., 2016).

The increased accessibility over the enhancer portion of the relocated II1 transgene in T-DOM suggested that HOX13 transcription factor may still be able to bind. We assessed this by performing CUT&RUN experiments using HOXA13 and HOXD13 antibodies on distal limb cells from embryos homozygous for the II1 transgene in T-DOM and wild type controls. To discriminate between reads from II1 in C-DOM or in T-DOM, only those reads containing sequences uniquely mapping to sequences outside the II1 element itself were considered (see methods). As expected, HOX13 proteins bound strongly to the native II1 element in C-DOM, with a peak over the 3’ end of the enhancer in controls (Figure 3A, left). At the analogous position of the II1 transgene in T-DOM, we observed a similar peak for both HOXA13 and HOXD13, indicating that HOX13 transcription factors were able to bind to the II1 enhancer element when inserted into the T-DOM (Figure 3A, right). However, the presence of these factors was not able to drive robust transcription of the nearby *LacZ* transgene in distal limb bud cells.

To confirm that the HOX13 peaks detected in the II1 enhancer element indeed corresponded to the presence of the expected HOX13 binding motif(s), we performed a motif search analysis (Heinz et al., 2010) for our CUT&RUN samples and for a previously reported dataset using ChIP-Seq (Sheth et al., 2016). In both datasets, we found motifs for HOX13 factors within the II1 element at four different positions. Three of these sites are closely clustered at the 3’ end of the enhancer element (Figure 3B, pink bars) and match the peak summit for HOXA13 and HOXD13 in both the native II1 enhancer environment in C-DOM and in the transgene in T-DOM (Figure 3A, grey columns). An additional motif was found within the II1 region, but locates outside the peak region (Figure 3B, asterisk). The three clustered motifs match previously reported HOX13 motifs (Desanlis et al., 2020; Sheth et al., 2016) and their position in relation to the position of CUT&RUN reads suggested that these sites are, in large part, responsible for the distal limb enhancer activity of the native II1 element in C-DOM.

### HOX13 binding sites are essential to II1 enhancer activity in distal limb buds

The presence of HOX13 factors bound to the II1 enhancer sequence when integrated into T-DOM suggested that they may be responsible for the weak remaining *LacZ* staining in some specific distal limb cells, even though this staining was distinct from the strong and general accumulation scored with the randomly integrated transgene (II1 TgN, compare Figure 1C and 2B). Instead, it could be caused by some other factor(s) taking advantage of the pioneer activity of HOX13 factors (Amândio et al., 2020; Desanlis et al., 2020). We verified this by testing the necessity of these three HOX13 binding sites in the II1 enhancer recombined into T-DOM. We implemented a CRISPR approach *in vivo* to delete the HOX13 binding sites in the II1 T-DOM transgene, by using guides to delete either two or three of the HOX13 binding sites (Figure 4A). Embryos hemizygous for the II1 enhancer in T-DOM were electroporated with CRISPR guides and Cas9 protein and the embryos were collected at E12.5 and stained for *LacZ*. Subsequently, the induced mutations were confirmed by Sanger sequencing (Table S2). While the *LacZ* staining in proximal limb cells was not affected by these mutations, in all cases where the binding sites were deleted, the remaining *LacZ* staining found in distal limb cells of II1 T-DOM transgenic embryos was completely ablated even after overstaining the samples (Figures 4B, Del TFBS and S4-1A).

**Figure 4.**
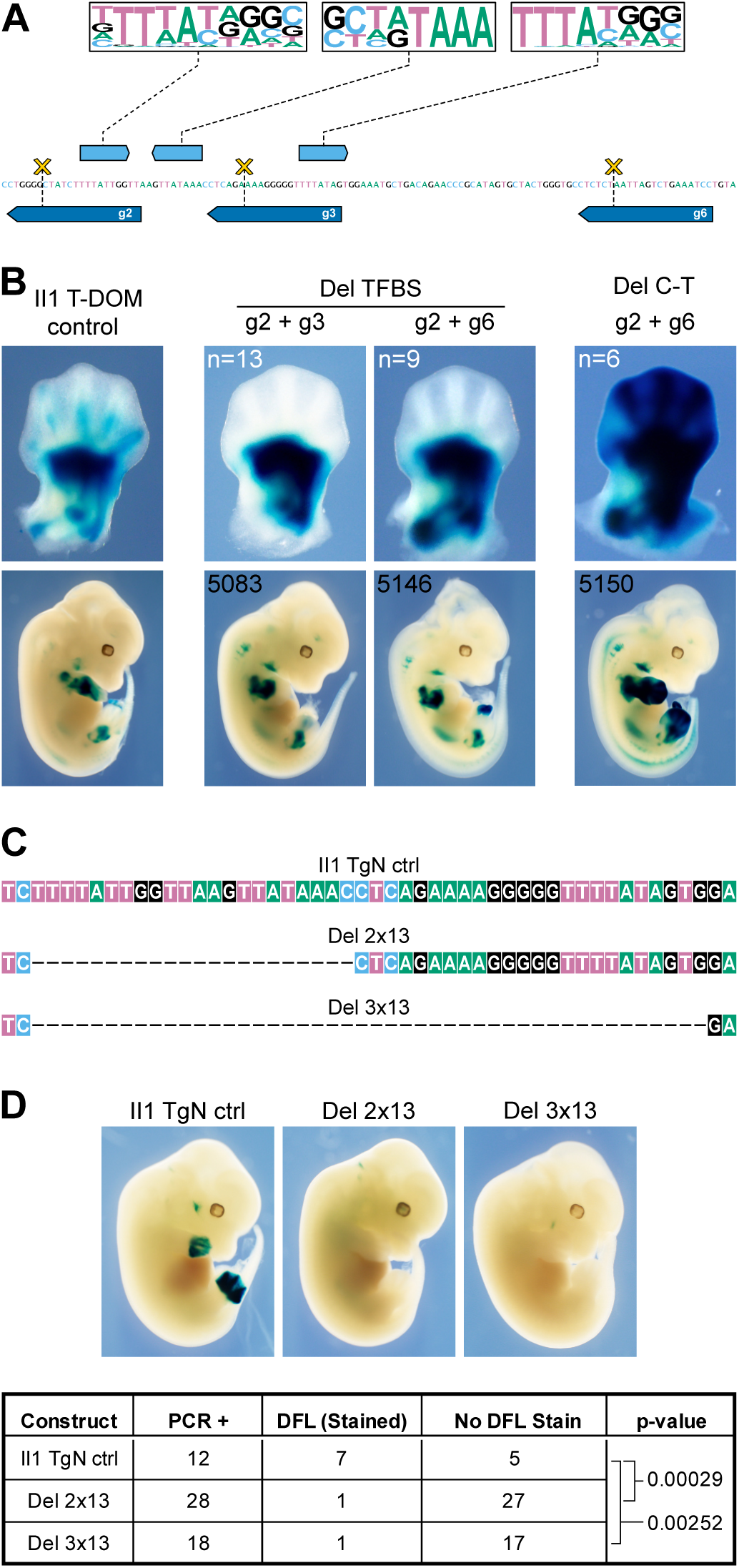
Deletions of HOX13 binding sites. **A**. On top are shown the HOX13 motifs in the II1 element, as extracted from the transcription factor binding (Figure 3B), with their positions indicated below (light blue boxes with orientations). The dark blue boxes below the DNA sequence indicate the positions and orientations of the CRISPR guides. Combinations of guides were used to generate small deletions (g2 and g3 or g2 and g6). The yellow crosses indicate the approximate cutting position of the Cas9. **B**. E12.5 F0 embryos stained for *LacZ*. The left panel is a II1 T-DOM limb showing the staining pattern with the recombined II1 enhancer (no CRISPR cutting), with weak staining in the distal limb, particularly in the digit mesenchyme. The two central panels are representative embryos (n indicated in upper left corners) with the indicated deletions (see Figure S4-1 for images of all embryos including these two #5083 and #5146). Embryos carrying either the g2+g3 or g2+g6 deletions lost all staining in distal limb cells. Several embryos showed strong distal limb staining (as shown in the right panel, see also Figure S4-1B). They contained a large deletion that extends from the native II1 element in C-DOM to the II1 transgenic element in T-DOM, due to the guide RNA sequences present at both sites (scheme in Figure S4-1C). Deletions in all embryos were sequenced (Table S2). Embryos with ambiguous sequencing results or mosaicism were not used. **C**. Sequence map of part of the II1 enhancer elements used to generate randomly integrated transgenic embryos. The top track is the wild type sequence around the three HOX13 binding sites for II1 used in control embryos (II1 TgN ctrl). The Del 2x13 sequence lacks the two centromeric HOX13 binding sites while the Del 3x13 sequence lacks all three HOX13 binding. **D**. On top are *LacZ* stained E12.5 transgenic embryos and the table on the bottom reports the number of embryos that were positive for the transgene by PCR (PCR+) followed but the number of embryos that stained in the distal forelimb (DFL) as well as the embryos with no staining in the DFL (No DFL Stain). The p-values are determined by Fisher’s exact test. This indicates that the 2x13 and 3x13 deletions are likely to be responsible for the loss of distal limb staining in these embryos compared with the II1 control embryos. The embryo on the left is a II1 control embryo containing the wild type II1 sequence. The embryos in the center and right are representatives of the staining (if any) obtained with the 2x13 and 3x13 constructs, with an absence of distal limb staining. Pictures of all embryos generated in this experiment containing some *LacZ* staining are in Figure S4-2, including the three shown here.

Of note, in several F0 embryos, we observed very strong proximal and distal limb *LacZ* expression (Figures 4B, S4-1B, Del C-T). When we sequenced the genomic DNA of these embryos, we found that they all contained a large deletion extending from the native II1 enhancer element within C-DOM up to the II1 transgene within the T-DOM region (approximately 1.2 Mb in length), due to the use of guide target sequences present in both copies of the II1 sequence (scheme in Figure S4-1C). In these embryos, the HOX13 binding sites are deleted and the *LacZ* transgene, formerly located within the T-DOM, was fused with a portion of the C-DOM (Figure S4-1C). In such a configuration, it is very likely that the enhancer remaining 5’ to the II1 site in C-DOM (island I) was then able to act on the transgene driving expression in the distal limb (Figures 4B and S4-1C), a regulatory influence obviously not permitted in the presence of an integral native T-DOM. Expression in proximal cells was controlled by those proximal enhancers located telomeric to the deletion breakpoint (Figure 2A, CS65 and PLEs).

As a control for the requirement of HOX13 factors for the function of the II1 enhancer sequence, we generated two variants of the II1 TgN transgene construct lacking either two (Del 2x13) or three (Del 3x13) HOX13 binding sites (Figure 4C, Table S1). We then injected these constructs into embryos to produce random integration events, and then stained for *LacZ*. In nearly all cases, neither the Del 2x13, nor the Del 3x13 variants were able to produce distal limb staining, when compared with the embryos containing the complete II1 enhancer sequence (Figure 4D). There were two exceptions to this: in one Del 2x13 embryo, we observed a clear distal limb staining although the proximal boundaries were very different than that seen in the normal II1 transgene (Figure S4-2, Del 2x13, yellow asterisk) and, in one Del 3x13 embryo, there was strong proximal limb staining and a portion of this staining extended into the posterior portion of the distal limb (approximately digit 5), yet most of the distal limb staining was absent (Figure S4-2, Del 3x13, yellow asterisk). In the remaining embryos carrying the transgene, twenty-seven embryos with the Del 2x13 transgene and seventeen embryos with the Del 3x13 transgene did not produce any distal limb staining (Figures 4D and S4-2).

As a final control experiment that integrates the two experiments above, we targeted the insertion of two additional variants of the II1 transgene into the same position of T-DOM as the II1 T-DOM 542 allele. In the first variant, we used the Del 3x13 transgene construction used to test the need for the three HOX13 binding sites when integrated at random locations throughout the genome (Figure 4C-D). When we stained embryos carrying this transgene, we again observed very strong proximal limb staining that matched the 542 allele, indicating that the transgene behaved as a T-DOM enhancer sensor in proximal limb cells (Figure S4-3A), but there was no staining in the distal limb. In the second variant we completely removed the II1 enhancer element from the transgene and inserted into the T-DOM. In this construction the *LacZ* staining pattern also produced strong proximal limb staining and no staining in the distal limb (Figure S4-3B).

Altogether, these results indicate that even in the presence of bound HOX13 proteins, which are normally the essential factors for its activation, the II1 enhancer sequence recombined within T-DOM was unable to express its full potential. Indeed, only a weak remnant of a transcriptional activity was scored in distal cells, at a late stage and low level, even though the reporter system could work at high efficiency in proximal cells. This suggested that, in distal limb cells, the surrounding chromatin context of T-DOM exerted a dominant effect to repress the activity that this sequence normally displays when positioned within C-DOM, even if the binding of HOX13 factors was still observed.

### A TAD-driven repression of distal enhancers in proximal limb cells?

In order to challenge this negative in-*cis* effect, we used CRISPR to delete portions of the T-DOM adjacent to the introduced enhancer transgene and then evaluated the effect of these deletions on transcription of the transgenic *LacZ* construct. Fertilized eggs hemizygous for the II1 T-DOM recombined enhancer allele were electroporated with CRISPR guides and Cas9 protein (Table S1). Since the DNA sequences targeted by guides are present on both the wild type and the II1 T-DOM alleles (Figure 5A, Table S1), we used genotyping PCR to screen for embryos carrying the expected deletion and used changes in *LacZ* staining to associate the deletion with the II1 T-DOM chromosome. As a control, we used littermate embryos that contained the same deletion but on the wild type (non-transgenic) chromosome.

**Figure 5.**
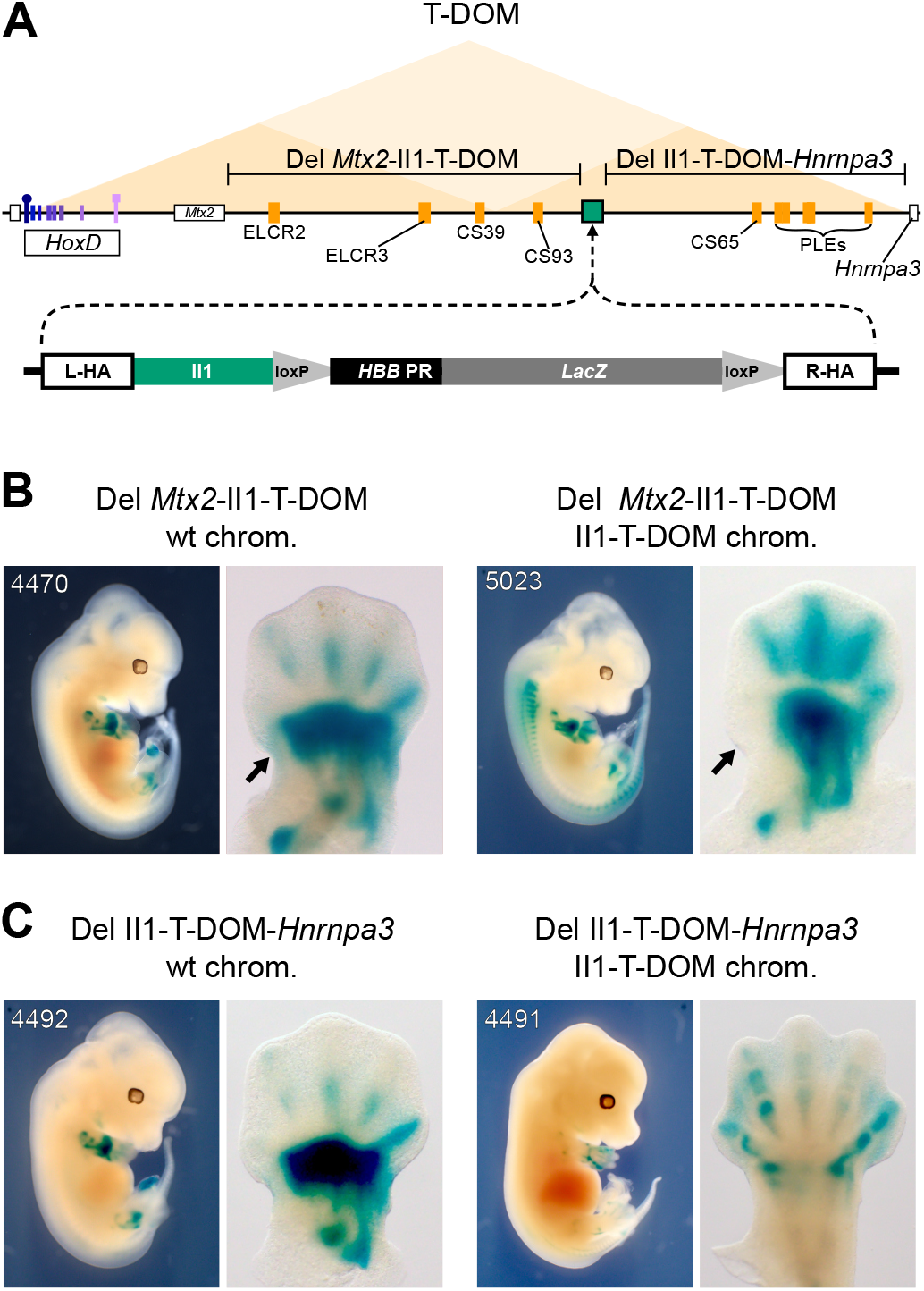
Inhibition of distal enhancer activity by the T-DOM chromatin environment. A. Schematic of T-DOM with the *Hoxd* gene cluster on the left (purple boxes), *Hoxd1* with a square and *Hoxd13* with a circle on top. The *Mtx2* gene is next to *Hoxd1* and *Hnrnpa3* gene is the small white box with black border on the right of T-DOM. The position of the II1 transgene insertion into T-DOM is indicated by a green rectangle with black border (mm10 chr2:75269597-75269616). Orange rectangles are known proximal limb enhancers. The two regions deleted by CRISPR are indicated above the genomic map (Del *Mtx2*-II1-T-DOM and Del II1-T-DOM-*Hnrnpa3*). **B**. Effect of deleting the centromeric portion of T-DOM (Del *Mtx2*-II1-T-DOM) on *LacZ* staining, with a light loss of staining in the proximal domain (black arrows) and a gain in the distal domain (see also Figure S5A). **C**. Effect of deleting the telomeric portion of T-DOM (Del II1-T-DOM-*Hnrnpa3*) on *LacZ* staining, with an almost complete loss of staining in the proximal domain (arrows) and no substantial impact on the distal domain (see also Figure S5B). The embryos shown here are also displayed in Figure S5 to show the complete series of stained embryos.

The first deletion extended from the 3’ end of the *Mtx2* gene, up to, but not including, the II1 enhancer element in the T-DOM (Figure 5A; Del *Mtx2*-II1-T-DOM). This deleted region of T-DOM contains the CS39 and CS93 proximal limb enhancers (Andrey et al., 2013; Yakushiji-Kaminatsui et al., 2018). In this deletion, we only scored a slight reduction in the extent of the *LacZ* staining in proximal limb cells (Figure 5B, black arrows), likely due to the removal of some proximal limb enhancers. In distal limb cells, there was a clear increase in the *LacZ* staining throughout the distal limb mesenchyme spanning almost the entire digital plate, as compared to the same deletion on the wild type chromosome (Figure 5B, compare left and right, Figure S5A).

The second deletion extended from the 3’ end of the *LacZ* transgene to the telomeric end of the T-DOM regulatory landscape (Figure 5A, Del II1-T-DOM-*Hnrnpa3*). This portion of T-DOM contains the CS65 and PLEs proximal limb enhancer elements (Andrey et al., 2013; Bolt et al., 2021). In this deletion we observed a severe loss of *LacZ* staining in proximal limb cells (Figure 5C, Figure S5B). In distal limb cells, there was a slight difference in the distribution of *LacZ* positive cells, yet no obvious increase in staining when compared to the control chromosome, unlike in the former deletion (Figure 5C, Figure S5B). These results showed that staining could be recovered when the centromeric flanking piece of T-DOM was removed and hence that this chromatin segment somehow exerted a robust repressive effect on the II1 transgene in distal limb bud cells.

### The II1 T-DOM transgene contacts the *HoxD* gene cluster

In distal limb bud cells at E12.5, strong chromatin contacts are detected between the II1 enhancer sequence (or a larger sequence including it) and the ‘posterior’ part of the *HoxD* cluster. Along with other C-DOM cis-regulatory regions, the II1-*Hox* cluster interactions collectively sustain activation of *Hoxd13* to *Hoxd11* in the digital plate (Montavon et al., 2011). Because of this, we wondered how this enhancer sequence would behave when relocated within the 3D chromatin space of the neighboring TAD. In other words, would it maintain its contacts with these ‘distal’ limb genes (*Hoxd13* to *Hoxd11*), not establish any contacts with the cluster at all, or would it adopt the interaction tropism of its new T-DOM neighborhood for the more ‘proximal’ limb genes (*Hoxd10*, *Hoxd9, Hoxd8*)? We performed Capture Hi-C (CHi-C) on E12.5 proximal and distal forelimb cells micro-dissected either from wild type embryos, or from embryos homozygous for the recombined II1 T-DOM transgene. The captured reads were mapped onto the mutant genome excluding all reads that would ambiguously map to both the II1 enhancer in C-DOM and in T-DOM, i.e., sequence reads that would not extend outside of the enhancer itself and hence could not be uniquely assigned to either one of the two sites.

In both the proximal and distal forelimb datasets, strong contacts were established between the II1 enhancer element within T-DOM and the *HoxD* gene cluster, as revealed by the subtraction of the mutant contact signal from the wild type (Figure 6A, black arrows; Figure S6A, black arrows). This gain of contacts between the recombined II1 enhancer and the *HoxD* cluster in both proximal and distal cells, were the only noticeable change induced by the presence of the transgene on the general chromatin configuration of the locus (Figure 6A). In order to evaluate these new contacts in greater detail, we plotted pairwise heatmaps between the *HoxD* cluster and II1 T-DOM, as well as H3K27me3, H3K27ac, and CTCF ChIP-seq datasets (Rodriguez-Carballo et al., 2017; Yakushiji-Kaminatsui et al., 2018).

**Figure 6.**
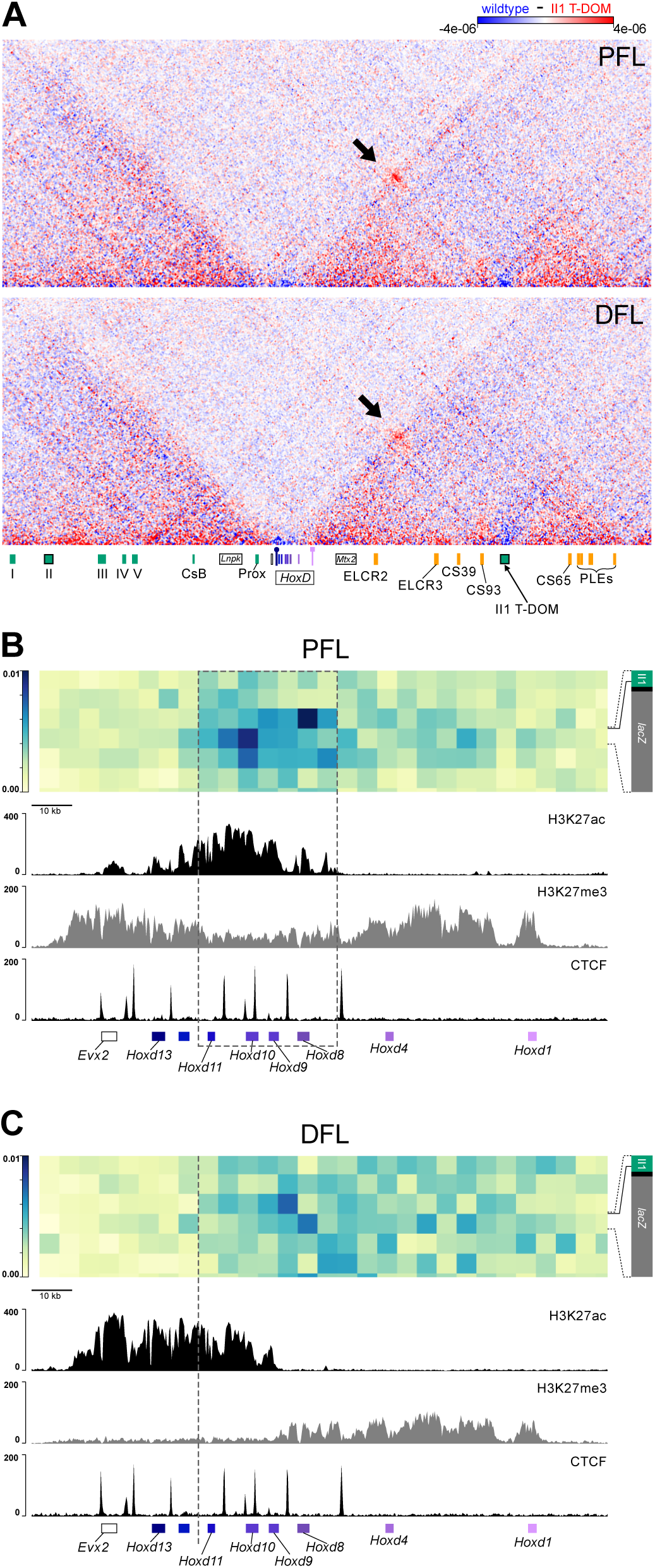
The II1 enhancer in T-DOM contacts the *Hoxd* gene cluster. **A.** Capture Hi-C maps at 5kb bin resolution over the entire *HoxD* locus (mm10 chr2:73950000-75655000), displayed as the subtraction of wild type signal (blue) from the II1 T-DOM 542 allele signal (red). The II1 enhancer T-DOM insertion site (arrow on the green rectangle at the bottom) produces increased contacts with the *Hoxd* gene cluster (black arrow pointing to the red bins). The contacts are established in both proximal forelimb (PFL) and distal forelimb (DFL) cells. **B.** Contacts between the *Hoxd* gene cluster (x-axis, genes are indicated below) and the region covering the II1 T-DOM reporter transgene (y-axis). The II1 T-DOM construct is schematized on the y-axis for clarity, with a solid black line indicating the boundary between bins. The II1 enhancer is shown in green, the *HBB* promoter is black, and the *LacZ* gene is in grey. The panels below are the H3K27ac, H3K27me3 (from (Yakushiji-Kaminatsui et al., 2018) and bound CTCF (from (Rodriguez-Carballo et al., 2017) from wild type PFL cells, aligned with the interaction matrix above. The strongest contacts are between the II1 T-DOM reporter transgene and the region around *Hoxd8* and *Hoxd10*, matching genes transcribed in proximal limb cells, indicated by a grey dashed box. **C.** Same as in B but using distal forelimb cells (DFL). The II1 T-DOM insert establishes more diffuse contacts, extending from *Hoxd1* to *Hoxd11* and stopping abruptly before those genes highly expressed in distal cells (*Hoxd12*, *Hoxd13*).

These alignments revealed clear differences between contacts in proximal and distal cells, in both their relative strengths and localization. In proximal cells, the II1 enhancer and the *HBB* promoter formed contacts mostly concentrated over the *Hoxd8* to *Hoxd10* region (Figure 6B, grey dashed box). However, even at this 5kb resolution, we were not able to resolve if the contacts were being mediated by the II1 transcription factor binding sites or the *HBB* promoter, because both portions of the transgene are within the same *DpnII* restriction fragment (Figure 6B, C). In distal limb bud cells, the contact dynamic was quite different. While the fragment containing the II1 enhancer and the *HBB* promoter continued to form the strongest contacts with the cluster, the region of highest contact had shifted from the 3’ end of *Hoxd8* to the 5’ end, and the robust contacts detected around *Hoxd10* in proximal cells had nearly disappeared (Figure 6C). Overall, the contacts were more evenly distributed over the region extending from *Hoxd1* to *Hoxd9*, but they were excluded from the *Hoxd12* to *Hoxd13* region (Figure 6C, grey dashed line).

The contacts between the II1 T-DOM transgene and the *HoxD* cluster changed between proximal and distal limb bud cells, exactly matching the distribution of active *versus* inactive chromatin in these two developmental contexts, respectively. In proximal limbs, the region of enriched contacts corresponded to a depletion in H3K27me3 marks and an enrichment in H3K27ac, which exactly matches the region of *HoxD* that is actively transcribed in proximal limb cells (Andrey et al., 2013; Tarchini and Duboule, 2006). In distal limb cells, the contacts became more evenly distributed and correlated with the presence of H3K27me3, stopping abruptly within the H3K27ac-positive portion of *HoxD* cluster, before reaching *Hoxd13* and *Hoxd12* (Figure 6C). Therefore, the II1 enhancer sequence, when recombined into T-DOM behaved spatially like a strong T-DOM proximal limb enhancers (Figure S6A), even though it had no intrinsic proximal limb specificity, as demonstrated by random transgenesis (Figure 1C, Figure 4D). Regardless of the context and the developmental time, when positioned into T-DOM, it never contacted the *Hoxd12* to *Hoxd13* region, which is the part of the cluster that this enhancer sequence normally contacts with the highest affinity in distal limb bud cells, nor did it influence in any way or in any cell type the chromatin structure that is normally found in T-DOM.

## DISCUSSION

The importance and status of enhancer sequences has evolved considerably since their discovery (Banerji et al., 1981). Initially described as short non-coding sequences that can increase the transcription rate of a target gene at a distance and regardless of orientation, they are now known to modulate gene expression in many different ways (Schaffner, 2015). Enhancers have a particular importance for genes with specialized expression patterns, producing transcription in specific cell types and tissues, at precise times and quantities, either during development or subsequently (Long et al., 2016). These sequences are thought to have evolved along with the emergence of novel body structures, recruiting genes already functional elsewhere, to accompany or trigger the formation of these novelties (Carroll et al., 2008). In vertebrates, where multi-functionality is common for genes having important functions during development, enhancers have accumulated in the vicinity of the transcription units forming regulatory landscapes (Spitz et al., 2003) that sometimes extend over several megabases (see (Bolt and Duboule, 2020). Within these landscapes, gene activation can be distributed across multiple enhancers (Marinic et al., 2013; Montavon et al., 2011) often leading to functional redundancy between them or to more complex interactions (Amândio et al., 2020; Osterwalder et al., 2018). Alternatively, enhancer sequences can be grouped together in a more compact manner, either to ensure a coordinated function, as exemplified by the *Globin* genes (Grosveld et al., 2021; Liu et al., 2018; Oudelaar et al., 2021), or to maximize transcription in a given cellular context such as the compact regulatory structure referred to as super enhancers (Blobel et al., 2021; Hnisz et al., 2013).

Enhancers are often embedded into Topologically Associating Domains (TADs), which are regions where certain DNA-DNA interactions are favored while adjacent regions are excluded from the interaction space (Dixon et al., 2012; Nora et al., 2012; Sexton et al., 2012). As illustrated at the *HoxD* locus, the genomic dimensions of TADs sometimes correspond to the extents of regulatory landscapes (Andrey et al., 2013), with TADs providing boundaries to the interaction space of enhancers within the three-dimensional organization of the genome. Consequently, TADs have been considered to be permissive structures augmenting enhancer and promoter interactions, while simultaneously providing borders that prevent enhancers from interacting with elements outside the TAD, and hence to regulate genes in an inappropriate manner (Lupianez et al., 2015, 2016). In this view, however, the enhancer sequence is considered as a regulatory element that can explore and act within the nuclear space somewhat freely unless TAD borders are present to frame its realm of action. Alternatively, there may be some loci where the function of an enhancer sequence can be subordinated to the global chromatin context of a given TAD thus introducing a level of regulatory control derived from a large chromatin domain rather than by individual DNA sequences (Andrey et al., 2013; Marinic et al., 2013; Rouco et al., 2021).

The transfer of enhancer II1 into T-DOM may illustrate such a case, where a potent and highly penetrant enhancer sequence is inhibited precisely in those cells where it normally functions, by placing it within a global environment that is not operational in these cells. In its normal location among C-DOM enhancers, the II1 sequence is bound by HOX13 factors (Beccari et al., 2016; Desanlis et al., 2020; Sheth et al., 2016), which are responsible for its strong activation potential as demonstrated here by the deletion of these sites in the transgenic context, which leads to the loss of *LacZ* staining in digit cells. When recombined into T-DOM, the II1 enhancer is silenced yet it still recruits HOX13 factors and hence its silencing cannot be attributed to the absence of the necessary activating factors. In fact, HOX13 factors are also bound to T-DOM whenever this TAD becomes inactive in distal limb cells and covered by H3K27me3 marks (Beccari et al., 2016), suggesting that the same factors may act in both a positive and a negative manner in different chromatin contexts.

One potential explanation to this observation is that HOX13 factors may function with more than one modality, depending on their context. On the one hand, these factors may bind to and activate an enhancer-reporter transgene in a sequence-specific manner, as many transcription factors do, leading to the pattern described herein and its absence in transgenes lacking the binding sites. On the other hand, both HOXD13 and HOXA13 proteins contain a large poly-alanine stretch (Bruneau et al., 2001), which may potentially drive the formation of phase-separated globules by co-condensing with transcriptional co-activators/co-repressors, as shown for these and other genes (Basu et al., 2020; Grosveld et al., 2021). It is thus possible that, due to their high content in bound HOX13 proteins, which contain stretches of poly-alanine (Bruneau et al., 2001), both C-DOM and T-DOM are used to form large transcription-hub condensate (Grosveld et al., 2021)(Amândio et al., 2020), leading to a positive transcriptional outcome for C-DOM in distal limb cells, and a negative outcome for T-DOM, within the same cells, due to the inclusion of additional co-factors that are specific to each domain (Karr et al., 2021). This latter explanatory framework would also account for why the deletion of the enhancer II1 *in vivo* had essentially no detectable effect upon transcription of target *Hoxd* genes in distal limbs, much like what was reported for C-DOM enhancers used for external genitals (Amândio et al., 2020) as well as in other comparable instances (Osterwalder et al., 2018). In both cases, removing a single component of the aggregate would not matter too much, whereas removing several related enhancers would then have a measurable impact.

This situation contrasts those where a single enhancer is responsible for target gene activation, the deletion of which usually seriously impairs the structure (see e.g. (Lettice et al., 2003; Shapiro et al., 2004). However, the former mechanism would not preclude the capacity for a single distal limb enhancer to trigger *LacZ* expression, as illustrated either by the weak staining detected when II1 was relocated into T-DOM, or when a large deletion brought the *LacZ* reporter close to Island 1, following the fusion between parts of C-DOM and T-DOM.

The II1 enhancer was selected because it is one of the strongest and most penetrant distal limb cells enhancer reported thus far (Lonfat et al., 2014). Yet it was silenced when introduced into T-DOM, along with several native T-DOM limb enhancers, which have a strong proximal specificity when inside T-DOM, while they can work efficiently in distal cells as well when randomly integrated as transgenes (Beccari et al., 2016). The repression of T-DOM in distal limb cells is reflected by the presence of large arrays of H3K27me3 marks (Andrey et al., 2013), and it is thus possible that the recombined II1 enhancer was included into this negative chromatin domain. The analysis of the II1 T-DOM 320 line (Figure 2C) suggests that this repression can be competed out by the presence of multiple copies of the enhancer-reporter construct, perhaps due to the formation of a particular sub-structure escaping the negative effect of T-DOM, or simply because of the accumulation of some transcription factors. This line was nevertheless not analyzed further due to technical difficulties associated with duplicated genomic sequences. We however conclude that special attention should be given to copy number when interpreting results from transgenic experiments.

The negative effect of T-DOM over the II1 enhancer construct *in-cis* was further suggested by our flanking deletions analyses. The (Del II1-T-DOM-*Hnrnpa3*) telomeric deletion did not substantially change the weak distal staining of the transgene, yet it severely reduced expression in the proximal domain, showing that the most potent proximal enhancers were located in the deleted interval. In contrast, the (Del *Mtx2*-II1-T-DOM) centromeric deletion consistently had the opposite effect, with only a slight reduction of the activity in the proximal limb domain while distal expression was re-activated, thus mapping the main T-DOM region carrying the repressive effect between the gene cluster and the integration site of the enhancer-reporter construct. However, it was not assessed whether this reactivated distal *LacZ* staining is due to the II1 enhancer itself or to the de-repression of other T-DOM enhancers, which would then act on the *HBB* promoter due to the in-*cis* proximity.

Finally, the recombined II1 construct was able to specifically and rather strongly contact the *HoxD* cluster, in both proximal (when T-DOM is active) and distal (when T-DOM is inactive) limb bud cells. In these two instances, however, the contacts were distributed differently. In proximal cells, contacts were established with the specific part of the cluster that is heavily transcribed, following the behavior of other T-DOM located enhancers (Andrey et al., 2013). In this case, these interactions were not directly driven by CTCF, for the integration site was selected at a distance from such sites. The enhancer was likely included into a global structure interacting with specific *Hoxd* promoters and possibly organized by CTCF sites within both the gene cluster and T-DOM (Rodríguez-Carballo et al., 2020). In distal cells, the interactions were less specific and likely reflected contacts between large H3K27me3 decorated chromatin segments (Noordermeer et al., 2011; Vieux-Rochas et al., 2015), as suggested by the absence of contacts with those genes heavily transcribed in distal cells. There again, the II1 enhancer behaved like its new neighboring proximal enhancers.

## CONCLUSION

Altogether, we conclude that in distal limb bud cells, the necessary decommissioning of all previously acting proximal enhancers is partly achieved -or secured-by a TAD-wide silencing mechanism, as illustrated by the appearance of large domains of H3K27me3. This mechanism is potent enough to prevent the expression of one of the strongest distal limb enhancers, after its recombination within T-DOM. This silencing is partly alleviated when a large piece of flanking chromatin is removed, suggesting the importance of neighboring sequences within the TAD to achieve this effect. When this distal enhancer was introduced into this ‘proximal TAD’, it behaved in all respects like its new neighbor proximal enhancers, thus illustrating the potential of chromatin domains, in some cases, to impose another level of coordinated regulation on top of enhancer sequence specificities.

## MATERIALS & METHODS

### Animal work

All experiments were approved and performed in compliance with the Swiss Law on Animal Protection (LPA) under license numbers GE 81/14 and VD2306.2 (to D.D.). All animals were kept in a continuous back cross with C57BL6 × CBA F1 hybrids. Sex of the embryos was not considered in this study. Mice were housed at the University of Geneva Sciences III animal colony with light cycle between 07:00 and 19:00 in the summer and 06:00 and 18:00 in winter, with ambient temperatures maintained between 22 and 23 °C and 45 and 55% humidity, the air was renewed 17 times per hour.

### Genotyping

When samples were to be used directly for experiments, a rapid protocol was implemented: Yolk sacs were collected and placed into 1.5 ml tubes containing Rapid Digestion Buffer (10mM EDTA pH8.0 and 0.1mM NaOH) then placed in a thermomixer at 95° for 10 min with shaking at 900 rpm. While the yolk sacs were incubating, the PCR master mix was prepared with Z-Taq (Takara R006B)(see Table S1 for genotyping primers) and aliquoted into PCR tubes. The tubes containing lysed yolk sacs were then placed on ice to cool briefly and quickly centrifuged at high speed. The lysate (1ul) was placed into the reaction tubes and cycled 32X (2s at 98°, 2s at 55°, 15s at 72°). 20ul of the PCR reaction was loaded onto a 1.5% agarose gel and electrophoresis was run at 120V for 10 minutes. Alternatively, when samples could be kept for some time, a more conventional genotyping protocol was applied; Tail Digestion Buffer (10mM Tris pH8.0, 25mM EDTA pH8.0, 100mM NaCl, 0.5% SDS) was added to each yolk sac or tail clipping at 250ul each along with 4ul Proteinase K at 20mg/ml (EuroBio GEXPRK01-15) and incubated overnight at 55°C. The samples were then incubated at 95° for 15 minutes to inactivate the Proteinase K and stored at -20°C until ready for genotyping. Genotyping primers (Table S1) were combined with Taq polymerase (Prospec ENZ-308) in 25ul reactions and cycled 2X with T_a_ = 64°C and then cycled 32X with T_a_ = 62°C.

### *LacZ* staining

Embryos were collected in ice cold 1X PBS in a 12-well plate. They were then fixed for 5 minutes at room temperature in freshly prepared 4% PFA with gentle shaking on a rocker plate. After fixing they were washed three times in washing solution (2mM MgCl2, 0.01% Sodium Deoxycholate, 0.02% Nonidet P40, and 1X PBS) for 20 minutes each at room temperature on a rocker plate. After approximately one hour of washing the wash solution was removed and replaced with staining solution (5mM Potassium Ferricynide, 5mM Potassium Ferrocynide, 2mM MgCl2 hexahydrate, 0.01% Sodium Deoxycholate, 0.02% Nonidet P40, 1mg/ml X-Gal, and 1X PBS). The plate was wrapped in aluminum foil and placed on a rocker plate over night at room temperature. The following morning the staining solution was removed and the embryos were washed three times in 1X PBS and then fixed in 4% PFA for long-term storage. Images of embryos were collected with an Olympus DP74 camera mounted on an Olympus MVX10 microscope using the Olympus cellSens Standard 2.1 software.

### Whole-mount *in situ* hybridization (WISH)

Embryos were collected at E12.5 and processed following a previously reported WISH procedure (Woltering et al., 2009). Briefly, embryos were fixed overnight in 4% PFA at 4°C. The following day they were washed and dehydrated through 3 washes in 100% methanol and then stored at -20°C until ready for processing. Each sample was prepared with Proteinase K (EuroBio GEXPRK01-15) at 1:1000 for 10 min. Hybridizations were performed at 69°C and post-hybridization washes were performed at 65°C. Staining was performed with BM-Purple (Roche 11442074001). All WISH were performed at on at least three biological replicates. Images of embryos were collected with an Olympus DP74 camera mounted on an Olympus MVX10 microscope using the Olympus cellSens Standard 2.1 software.

### RT-qPCR

Embryos were isolated from the uterus and placed into 1x DEPC-PBS on ice. The yolksacs were collected for genotyping. The embryos were transferred into fresh 1x DEPC-PBS and the distal limb portion was excised, placed into RNALater (ThermoFisher AM7020), and stored at -80°C until processing. Batches of samples were processed in parallel to collect RNA with Qiagen RNEasy extraction kits (Qiagen 74034). After isolating total RNA, first strand cDNA was produced with SuperScript III VILO (ThermoFischer 11754-050) using approximately 500ng of total RNA input. cDNA was amplified with Promega GoTaq 2x SYBR Mix and quantified on a BioRad CFX96 Real Time System. Expression levels were determined by dCt (GOI - Tbp) and normalized to one for each condition by dividing each dCT by the mean dCT for each wild type set. Table S1 contains the primer sequences used for quantification. Box plots for expression changes and two-tailed unequal variance t-tests were produced in DataGraph 4.6.1.

### CUT&RUN

Embryos were collected in ice-cold 1X PBS and yolk sacs were processed according to the rapid genotyping protocol described above. Embryos with the correct genotype were transferred to fresh PBS and dissected. The dissected tissue samples were transferred into 1X PBS containing 10% FCS and then digested with collagenase (see ATAC-Seq protocol below). For the HOXD13 and HOXA13 CUT&RUN, pools of cells from individual embryos were processed. All samples were processed according to the CUT&RUN protocol (Skene et al., 2018) using a final concentration of 0.02% digitonin (Apollo APOBID3301). Cells were incubated with 0.5 ug/100ul of anti-HOXD13 antibody (Abcam ab19866), 0.5ug/100ul of anti-HOXA13 (Abcam Ab106503) in Digitonin Wash Buffer at 4°C. The pA-MNase was kindly provided by the Henikoff lab (Batch #6) and added at 0.5ul/100ul in Digitonin Wash Buffer. Cells were digested in Low Calcium Buffer and released for 30 minutes at 37°C. Sequencing libraries were prepared with KAPA HyperPrep reagents (07962347001) with 2.5ul of adaptors at 0.3uM and ligated for 1 hour at 20°C. The DNA was amplified for fourteen cycles. Post-amplified DNA was cleaned and size selected using 1:1 ratio of DNA:Ampure SPRI beads (A63881) followed by an additional 1:1 wash and size selection with HXB. HXB is equal parts 40% PEG8000 (Fisher FIBBP233) and 5M NaCl.

CUT and RUN libraries were sequenced paired-end on a HiSeq4000 sequencer and processed as in (Bolt et al., 2021), mapped either on mm10 or on the II1TDOM-542 mutant genome. The E11.5 whole-forelimb HOXA13 and HOXD13 ChIP-Seq datasets (SRR3498934 of GSM2151013 and SRR3498935 of GSM2151014) as well as E12.5 distal and proximal forelimb H3K27Ac and CTCF ChIP-Seq datasets (SRR5855214 of GSM2713703, SRR5855215 of GSM2713704, SRR5855220 of GSM2713707 and SRR5855221 of GSM2713708) were processed similarly to what has been previously published (Beccari et al., 2021). Adapter sequences and bad quality bases were removed with Cutadapt [Martin et al. https://doi.org/10.14806/ej.17.1.200.] version 1.16 with options -a GATCGGAAGAGCACACGTCTGAACTCCAGTCAC -A GATCGGAAGAGCGTCGTGTAGGGAAAGAGTGTAGATCTCGGTGGTCGCCGTATC ATT -q 30 -m 15 (−A being used only in PE data sets). Reads were mapped with bowtie (Langmead and Salzberg, 2012) 2.4.1 with default parameters on mm10. Alignments with a mapping quality below 30, as well as discordant pairs for PE datasets, were discarded with samtools view version 1.8 <u>(Li, 2011; Li et al., 2009)</u>. Coverage and peak calling were computed by macs2 (Zhang et al., 2008) version 2.1.1.20160309 with options --bdg --call-summits -- gsize ’1870000000’, and -f BAMPE for PE. The HOX13 motifs where identified with findMotifsGenome.pl from the Homer tool suite (Heinz et al., 2010) using the narrowPeak of HOXA13 and HOXD13 ChIP as well as the third replicate of the HOXA13 and HOXD13 CUT and RUN with the option -size 50. The best motif of each of these four datasets was used to scan the sequence of the II1 enhancer. Four motifs were identified, for the three displayed in Figure 2B, the logo of the motif giving the best score is shown.

### ATAC-Seq

Samples used for ATAC-Seq were processed following the original protocol. Cells were collected in 1X PBS on ice and yolk sacs were collected for each sample. Embryos were rapidly genotyped (see above) and those with the correct genotype were transferred to fresh 1X PBS and dissected. Tissue samples were transferred into 300ul 1X PBS containing 10% FCS on ice until ready for processing. To each sample, 8ul of collagenase (at 50mg/ml, Sigma C9697) was added and tubes were placed in a Thermomixer at 37° with shaking at 900rpm for approximately 5 minutes or until the samples were completely disaggregated. The samples were then placed into a centrifuge at 4°C and centrifuged at 500xg for 5 minutes. The supernatant was removed and cells were gently resuspended in ice-cold 1X PBS. The cells in each sample were counted with a Countess (ThermoFisher) using Trypan Blue and checked for viability >90%. The volume needed to contain was 50,000 cells was determined and that volume was transferred to a new tube and centrifuged at 4°C at 500xg for 5 minutes. The supernatant was removed and the cell pellet was gently resuspended in lysis buffer then immediately centrifuged at 500g for 10 minutes at 4°C. The lysis buffer was removed and the cell pellet was gently resuspended in 50ul tagmentation mix (Nextera FC-121-1030) and then incubated at 37° in a thermomixer at 300rpm for 30 minutes. The samples were then mixed with Buffer QG from the Qiagen MinElute PCR Purification kit (28004) and processed according to that protocol, and then eluted from the column in 11ul EB. Samples were stored at -20°C until ready for library preparation. For library preparations, the samples were amplified with Nextera Index Primers (FC-121-1011) using NEBNext High-Fidelity 2x PCR Master Mix (M0541) and cycled 11 times. After PCR the reactions were cleaned first with the Qiagen MinElute PCR Purification Kit and then with AMPure XP beads (A63881) at a ratio of 1.8:1.0 followed by elution with 15ul EB.

Adapter sequences and bad quality bases were removed with Cutadapt (Martin, 2011) version 1.16 with options -a CTGTCTCTTATACACATCTCCGAGCCCACGAGAC -A CTGTCTCTTATACACATCTGACGCTGCCGACGA -q 30 -m 15. Reads were mapped with bowtie (Langmead et al., 2009) 2.4.1 with parameters -I 0 -X 1000 --fr --dovetail --very-sensitive on mm10 or on the II1TDOM-542 mutant genome. Alignments with a mapping quality below 30, discordant pairs, and reads mapping to the mitochondria, were discarded with bamtools version 2.4.0 [https://github.com/pezmaster31/bamtools]. PCR duplicates were removed with Picard[http://broadinstitute.github.io/picard/index.html] version 2.18.2 before the BAM to BED conversion with bedtools <u>(Quinlan, 2014)</u> version 2.30.0. Coverage and peak calling were computed by macs2 (Zhang et al., 2008) version 2.1.1.20160309 with options -- format BED --gsize 1870000000 --call-summits --keep-dup all --bdg --nomodel --extsize 200 --shift -100.

### Capture Hi-C

Samples used in the Capture Hi-C were identified by PCR screening embryos at E12.5 as described above. Collagenase treated samples were cross-linked with 1% formaldehyde (ThermoFisher 28908) for 10 minutes at room temperature and stored at -80° until further processing. The SureSelectXT RNA probe design used for capturing DNA was done using the SureDesign online tool by Agilent. Probes cover the region mm9 chr2:72240000-76840000 producing 2x coverage, with moderately stringent masking and balanced boosting. Capture and Hi-C were performed as previously reported. Sequenced DNA fragments were processed as previously reported but the mapping was performed on a mutant genome reconstructed from minion and Sanger sequencing (see below). A custom R (www.r-project.org) script based on the SeqinR package (Charif and Lobry, 2007) was used to construct a FASTA file for the mutant chromosome 2 from the wild-type sequence and the exact position and sequence of breakpoints.

Subtraction of matrices was performed with HiCExplorer (Ramirez et al., 2016; Wolff et al., 2020, 2011) version 3.6. Heatmaps were plotted using a custom version (Lopez-Delisle et al., 2021) of pyGenomeTracks (Ramírez et al., 2018) based on 3.6.

### II1 TgN Cloning and transgenesis

The II1 enhancer sequence (mm10 chr2:74075305-74075850) was amplified from the fosmid clone WI1-109P4 using primers 001 and 002 (Table S1). The 001 primer contains a *XhoI* site and LoxP sequence followed by sequence to the II1 enhancer. The 002 primer contains at its 5’ end a *HindIII* site. This PCR product was gel purified with Qiagen Gel Extraction Kit (28704). The PCR fragment and the pSKlacZ reporter construct were digested with *XhoI* and *HindIII* and ligated together with Promega 2X Rapid Ligation kit (C6711) to produce pSK-II1-LoxP-*LacZ* construct. This vector (15ug) was cut with *XbaI* and *XhoI* to release the enhancer-reporter construct. The digest was separated on a 0.7% agarose gel for 90 minutes. The 4190bp fragment was excised from the gel and purified with Qiagen Gel Extraction Kit (28704) and eluted in 30ul EB followed by phenol-chloroform extraction and ethanol precipitation and then the pellet was dissolved in 30ul TE (5mM Tris pH7.5, 0.5mM EDTA pH8.0). DNA was injected at 3ng/ul into pronuclei. Five founder animals were identified carrying the transgene by PCR. Four male founders with the transgene were put into cross with wild type females and embryos were collected at E12.5 to test for *LacZ* staining. All four male founder lines (*LacZ*/40, 41, 44, and 46) produced distal limb staining but *LacZ*/40 was chosen for amplification of the breeding line due to high transmission of the transgene.

### II1 T-DOM targeted insertion

The II1 TgN transgenic construct, outlined above, was used for the targeted insertion but homology arms (HA) were attached (Left HA: mm10 chr2:75268556-75269591, Right HA: mm10 chr2:75269617-75270665). The cloning vector was linearized with *KpnI*, separated on a 0.7% agarose gel for 90 minutes at 90 volts, then the 9.0kb band was extracted and purified two times with Qiagen Gel Extraction kit (28704). The DNA was quantified by Qubit dsDNA and diluted to 5ng/ul with IDTE (11-05-01-05). Wildtype fertilized eggs were injected with the construct and the supercoiled pX330 (Addgene #42230) expression vector containing the sgRNA sequence (r4g9: mm10 chr2:75269597-75269616). Animals were genotyped (Table S1) to identify founders, and then sequenced with minION (below).

### minION Sequencing

Long-read sequencing was performed on the II1 T-DOM alleles (II1 T-DOM 320 and II1 T-DOM 542) following the nCATS protocol with minor changes (Gilpatrick et al., 2020). Yolk sacs were isolated from embryos containing the II1 T-DOM transgene and digested with Tail Digestion Buffer (see above) and Proteinase K overnight at 55°C with no shaking. The following day the samples were incubated at 95°C for 10 minutes to inactivate the Proteinase K followed by ethanol precipitation and eluted in 200ul of 10mM Tris pH7.5. CRISPR guides (Table S1) were designed in CHOPCHOP v3.0 (Labun et al., 2019) and synthesized as Alt-R RNAs by IDT. CRISPR crRNAs were duplexed with tracrRNAs according to the IDT protocol (Alt-R CRISPR-Cas9 System: In vitro cleavage of target DNA with ribonucleoprotein complex, version 2.2). Two master mixes of guide RNAs and Cas9 protein (1081059) were prepared (see Supplementary Figure 3 for sequence and cutting locations with the locus map), containing either SCS-12 and SCS-13 or SCS-14. The gDNA (9ug) from the yolk sac was dephosphorylated with NEB Quick CIP (M0510) for 10 minutes at 37°C followed by 2 minutes at 80°C to inactivate the CIP. The gDNA was split into two equal pools and each pool was then combined with the guideRNP master mixes to cut the gDNA for 30 minutes at 37°C followed by 5 minutes at 72°C. The samples were then A-Tailed and AMX adaptors were ligated (Oxford Nanopore SQK-LSK109). The reactions were size selected with 0.3X Ampure SPRI beads (A63881) followed by two washes with Long Fragment Buffer and then eluted for 30 minutes in 15ul EB. The DNA libraries were then prepared according to the Oxford Nanopore protocol for sequencing on a minION (ENR_9084_v109_revP_04Dec2018). The sequencing ran for approximately 24 hours and was stopped for processing after all nanopores were depleted.

Bases were called from the fast5 files using Guppy base-caller (Oxford Nanopore Technologies) for CPU version 5.0.16+b9fcd7b. Reads were mapped on mm10 with minimap2 (Li, 2018) version 2.15 with parameter -ax map-ont. Only primary alignments were kept with samtools view version 1.10 (Li, 2011; Li et al., 2009) and reads mapping to II1 (mm10:chr2:74073413-74076528) or to the insertion region (mm10:chr2:75262998-75286118) were further analyzed. Read sequences were compared to the wild-type genome, the expected mutant genome as well as the sequence of the cloning vector using a Perl script as in (Schmidl et al., 2015) with the following modification: 20 bp of the MinION reads were tested against the reference for 5 bp-sliding windows and only 20-mers completely identical to unique 20-mers in the reference were kept. The output was then processed in R (www.r-project.org) to display dot plots. The in-depth analysis of reads for allele 320 allowed to propose a configuration that would match all reads containing in total four times the II1-*LacZ* construct.

### Del II1 TFBS TgN cloning and transgenesis

The enhancer constructs with mutations (Figure 4C) in the transcription factor binding sites were constructed *in silico* and synthesized by TWIST Bioscience (San Francisco, CA). The enhancer sequences are available in Table S1. The enhancer sequences were synthesized with *BglII* and *AgeI* restriction sites at the 5’ and 3’ ends respectively. The mutant enhancer sequences (Del 2x13 and Del 3x13) were restriction digested along with pSKlacZ and ligated together with Promega 2X Rapid Ligation kit to produce pSK-II1Del2X13lacZ and pSK-II1Del3X13lacZ. The enhancer-reporter fragments were released from the vector with *BglII* and *XhoI* and purified as above. Pro-nuclear injections were performed by the transgenic platform of the University of Geneva, medical school (CMU). Embryos were collected at approximately E12.5 and stained for *LacZ*.

### Deletions of transcription factor binding sites within T-DOM *in vivo*

Guide sequences were selected from the UCSC mm10 genome browser track CRISPR/Cas9 - NGG Targets. The crRNA Alt-R guides were synthesized by IDT. Males homozygous for the II1 T-DOM 542 allele were crossed with super-ovulated wild type females (BL6XCBA-F1) and fertilized eggs were collected. The embryos were electroporated with CRISPR guides (12ug of each guide) and TrueCut Cas9 v2 protein (Thermo Fisher A36497) with a NEPA21 (NEPA GENE Co. Ltd, Chiba, Japan) and then reimplanted into surrogate females. Embryos were collected at E12.5. Yolk sacs were digested and the II1:*HBB:LacZ* transgene was PCR amplified and Sanger sequenced to identify transgenic embryos containing the mutagenized enhancer element. Embryos that were mosaic for the mutation were not included in this analysis. The embryos were *LacZ* stained (see above) at 37° for 16 hours, washed in PBS and post-fixed, then stored in 70% ethanol for photographing.

### Generation of Del *Mtx2*-II1-T-DOM and Del II1-T-DOM-*Hnrnpa3* alleles *in vivo*

The same methodology was used for this experiment as in the Del II1 T-DOM TFBS experiment above. The sequences for CRISPR guides used in this experiment are listed in Table S1. The embryos were genotyped for the presence of the deletion using primers in Table S1. Embryos with ambiguous genotyping results were not used in these results.

### Targeted insertion of Del 3x13 and *HBB*:*LacZ* transgene in T-DOM of ESCs

The two targeted insertions of control transgenes into T-DOM, genetic editing, and cellular culture were performed as previously reported (Figure S4-3) (Andrey and Spielmann, 2017; Kraft et al., 2015). The guide sequence (r4g9, Table S1) was cloned into pX459 vector from Addgene (#62988) and 8ug of the vector was used for mESC transfection. The pX459 vector was co-transfected with 4ug of the vector containing one of the two transgene cassettes. The two transgene cassettes (Del 3x13 and HBB:lacZ) contain the same vector backbone with the HBB promoter, lacZ gene, and SV40 polyA signal and homology arms used to target the r4 region of T-DOM (mm10 chr2:75269597-75269616). This is the same insertion site as the II1 T-DOM 542 and 320 alleles. The Del 3x13 contains the II1 enhancer element but with the three HOX13 binding sites removed (see Figure 4C, D). The HBB:lacZ transgene is the same but does not contain any portion of the II1 enhancer element so that it is strictly an enhancer sensor in the T-DOM. These constructs were co-transected into G4 mESCs obtained from the Nagy laboratory (George et al., 2007). After genotyping to confirm the insertion, the desired mESCs were thawed, seeded on male and female CD1 feeders and grown for 2 days before the aggregation procedure. ESCs were then aggregated with tetraploid (c57bl6J x B6D2F1) morula-stage embryos and let developed until blastula prior to transfer into CD1 foster females by the transgenic mouse platform at the University of Geneva Medical School (Artus and Hadjantonakis, 2011).

## ACKNOWLEDGEMENTS

We thank Jozsef Zakany for his help in an initial phase of this work and other colleagues from the Duboule laboratories for discussions This work was supported in part using the resources and services of the Gene Expression Research Core Facility (GECF) at the School of Life Sciences of EPFL and the transgenic platform at the medical school, University of Geneva.

## ETHICS APPROVAL

All experiments involving animals were performed in agreement with the Swiss Law on Animal Protection (LPA), under license No. GE 81/14 (to DD).

## DATA AVAILABILITY

All raw and processed datasets are available in the Gene Expression Omnibus (GEO) repository under accession number GSE194114. All scripts necessary to reproduce figures from raw data are available at https://github.com/lldelisle/scriptsForBoltEtAl2022

## AUTHOR CONTRIBUTIONS

C.C.B.: Designed and conducted experiments, analyzed datasets, formalized results and wrote the paper.

L.L-D. : Analyzed and evaluated the statistical significance of datasets. Wrote the paper. B.M.: Produced and maintained all of the mouse lines.

A.H.: Performed the Del 2x13 and 3x13 experiment

A.R.: Performed the Del 3x13 T-DOM experiment

G.A.: Designed and performed the *HBB*:*LacZ* T-DOM experiment

D.D.: Designed experiments, transported mice, dissected some limb buds and wrote the paper.

---

**Supplementary Figure 1.**
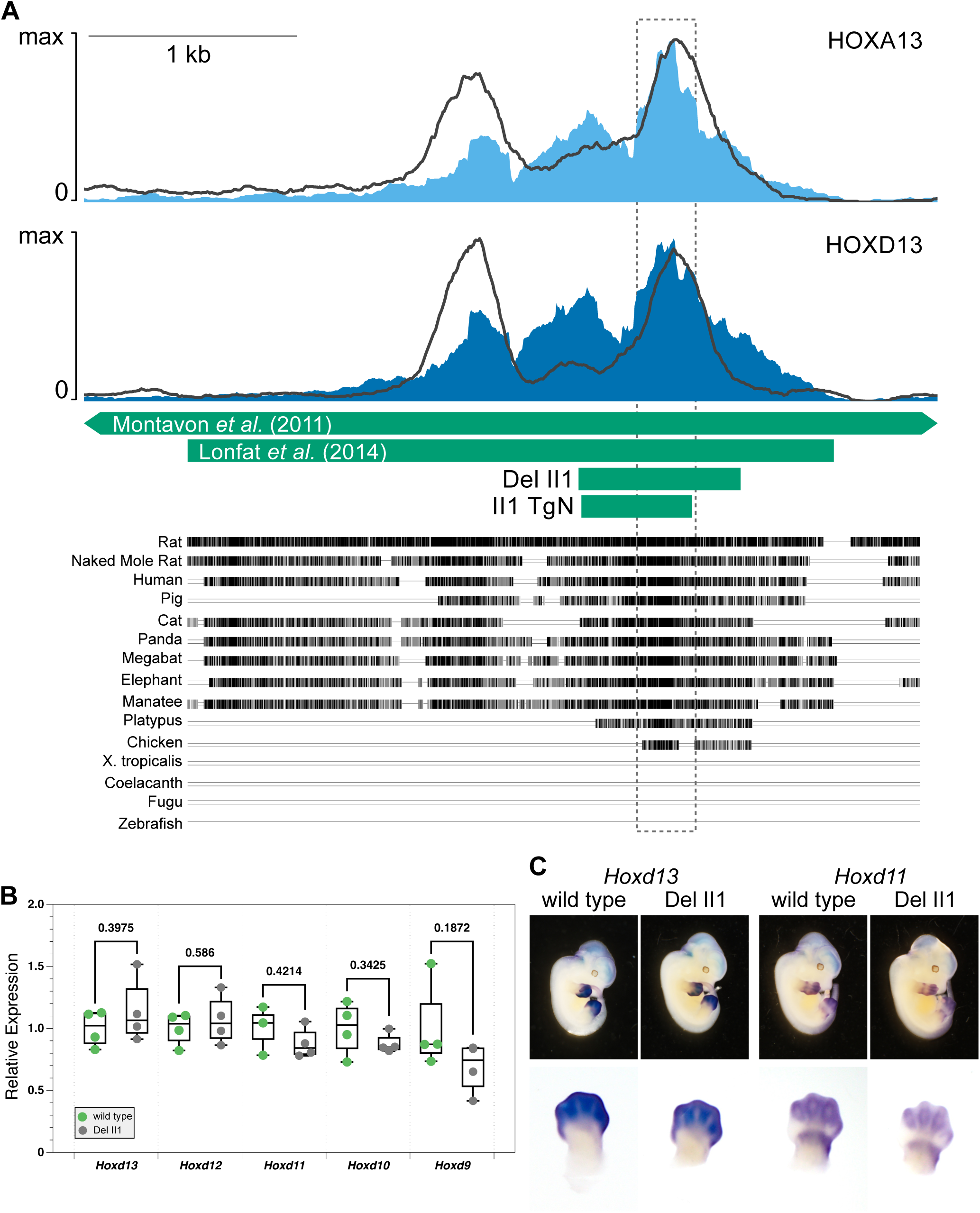
Deletion of the II1 enhancer does not alter *Hoxd* gene expression in the distal limb bud cells. **A**. Magnified representation of the Island II1 enhancer element (mm10 chr2:74072912-74077028, (Montavon et al., 2011). HOXA13 (top) and HOXD13 (bottom) E12.5 CUT&RUN profiles are indicated by light or dark blue, whereas HOXA13 and HOXD13 E11.5 forelimb ChIP-seq profiles (Sheth et al., 2016) are shown as dark grey lines super-imposed for comparison. The DNA sequence conservation track MultiZ from UCSC (mm10) is shown below, with the most conserved region indicated with a bracket. **B.** RTqPCR for multiple *Hoxd* genes using distal forelimb bud cells from E12.5 wild type and homozygous Del II1 mutant specimen. P-value is indicated above, no significant difference in expression was detected for any *Hoxd* genes in the absence of the II1 enhancer (two-tailed unequal variance t-test). **C**. Whole-mount *in situ* hybridizations for *Hoxd13* and *Hoxd11* in embryos homozygous for the II1 enhancer deletion also show no change in their expression domains when this enhancer is deleted from C-DOM.

**Supplementary Figure 2.**
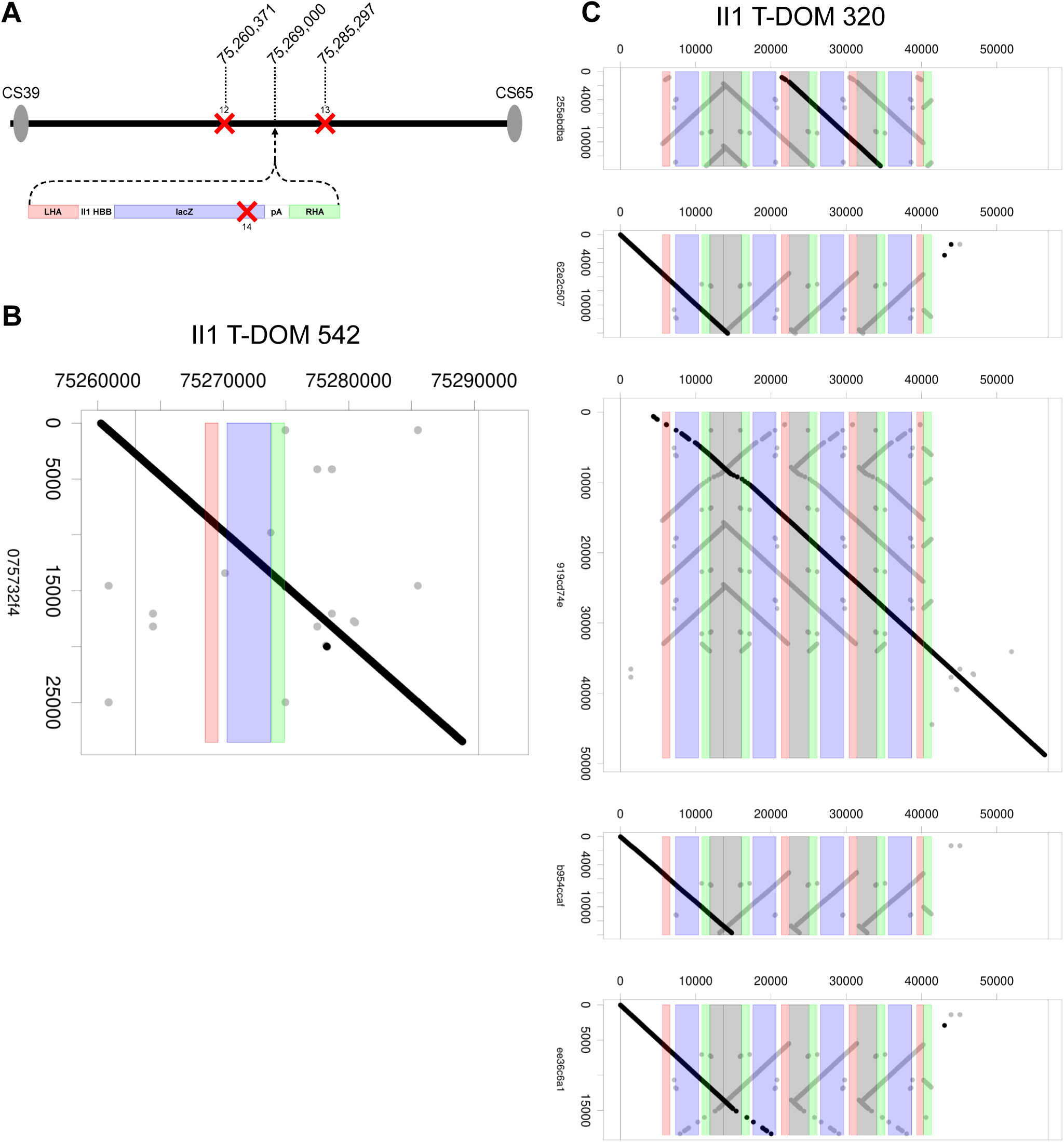
Long-read sequencing of the genomic structure after targeted insertion. **A**. Map of the T-DOM after homologous recombination of the II1:*HBB:LacZ* enhancer-reporter transgene, indicating the location of CRISPR guides used in the nCATS protocol for enrichment of sequencing reads (Gilpatrick et al., 2020). The red crosses indicate the location of the CRISPR cutting guide (see Supplementary Table S1). Two guides were used outside the II1 transgene (SCS12 and 13) and one guide within the II1 transgene (SCS14). Below the map of the CRISPR cutting position is a map of the transgene construct. The colors indicated for different portions of the construct match to the sequencing alignments below. **B**. Dotplot maps of sequencing reads recovered from the II1 T-DOM 542 allele and showing a clear one-copy recombination at the expected site. The x-axis is the position along the mutant II1 T-DOM 542 chromosome and the values on the y-axis represent the base pair position of the transgene. Each circle represents a 20bp alignment (see methods) so multiple adjacent 20bp matching reads appear as a line. The best matching read is drawn in black, while shorter reads are drawn in grey. **C**. Dotplot maps of five sequencing reads recovered from the II1 T-DOM 320 allele, showing insertion of multiple copies.

**Supplementary Figure 4-1.**
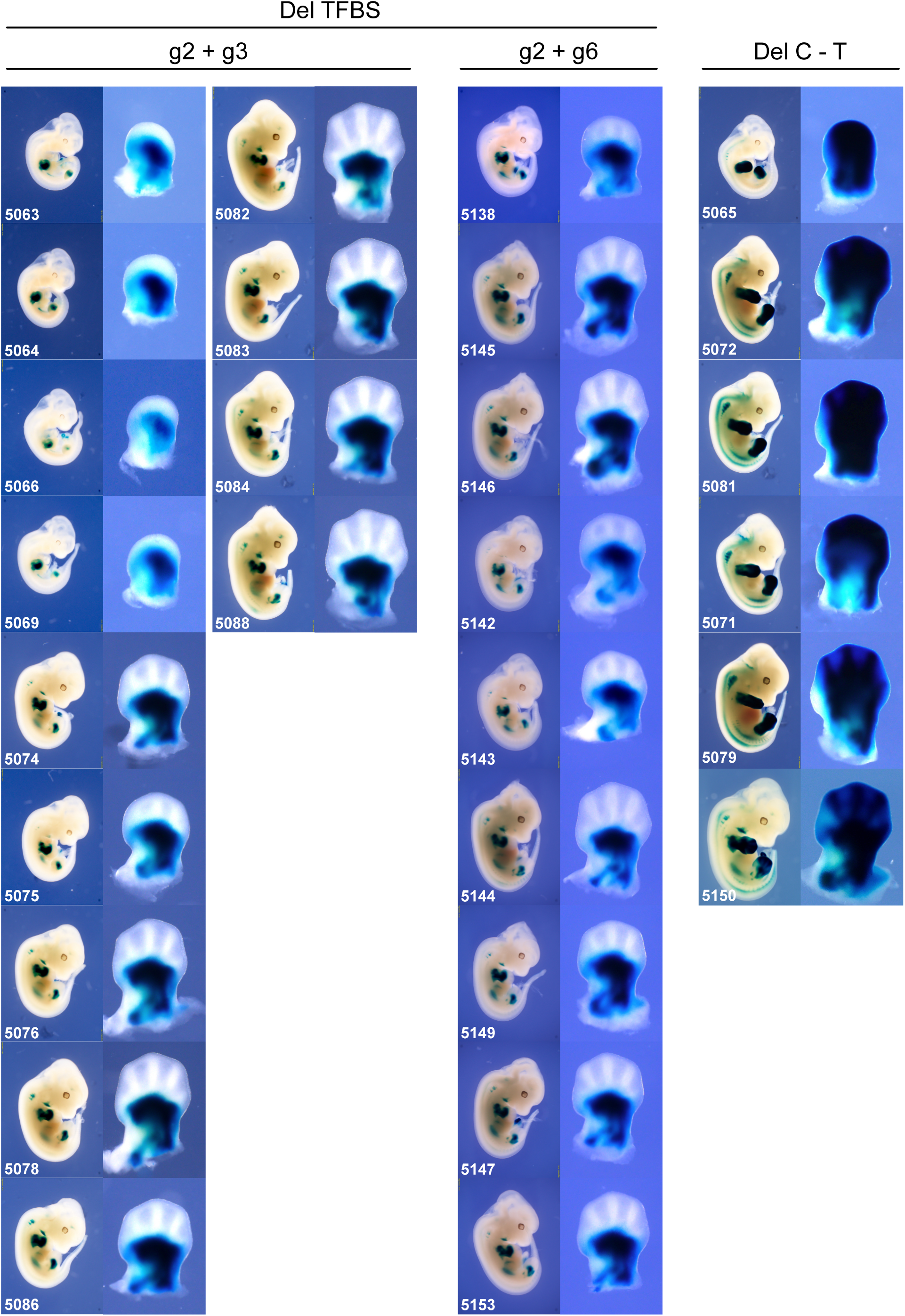
Photos of all embryos containing the deletions of the HOX13 binding sites present in the II1 enhancer-reporter construct targeted into T-DOM. Individual embryos are numbered. Embryos #5083 and #5146 are already shown in Figure 4B, but are reproduced here for easier comparison. The g2 + g3 and g2 + g6 deletions removed only binding sites within the II1 enhancer. **B**. Staining of embryos containing the Del C-T deletion. These embryos are positive in both proximal and distal limb bud cells. Embryo #5150 is already shown in Figure 4B but is added here for comparison. **C**. Schematic of the Del C-T deletion. The Del C-T created a large deletion fusing parts of T-DOM and C-DOM due to the presence of the sequence targeted by the guides RNAs on both native (C-DOM) and transgenic (T-DOM). After deletion, the II1 *LacZ* reporter transgene is flanked by a centromeric distal limb enhancer (green) and several telomeric proximal enhancers (CS65, PLEs, orange), thus accounting for its expression in both limb domains.

**Supplementary Figure 4-2.**
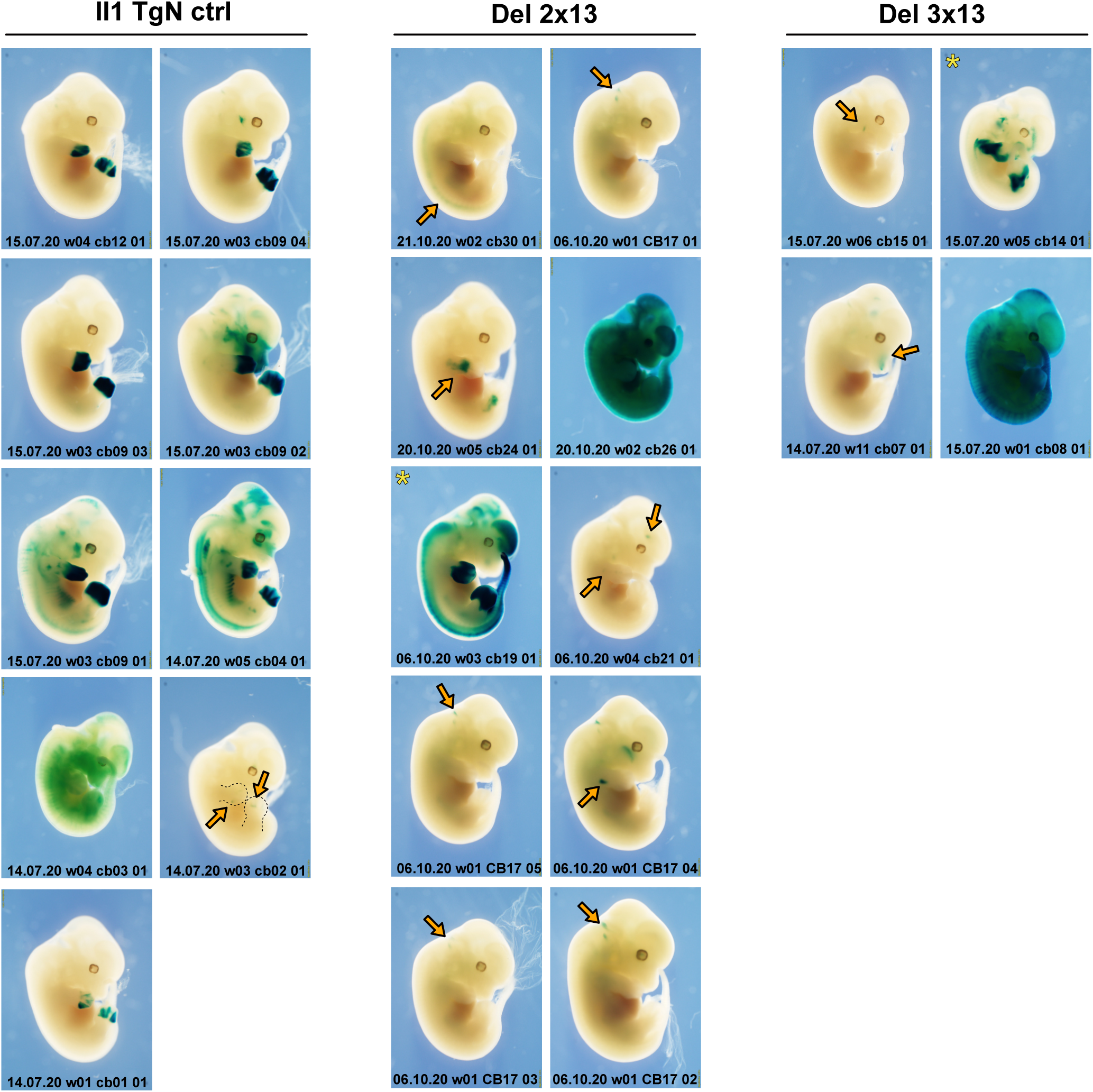
Photos of randomly integrated transgenic embryos stained for *LacZ* expression when HOX13 binding sites were deleted from the II1 enhancer. The left two columns (II1 TgN ctrl) are control embryos containing the entire II1 enhancer sequence. The two columns in the center (Del 2x13) show stained embryos containing the II1 enhancer element lacking the two centromeric HOX13 binding sites (see Figure 4A, C). The two columns in the right (Del 3x13) show stained embryos lacking all three HOX13 binding sites. The orange arrows indicated the location of *LacZ* staining when it is not detected in the distal limb buds. The unique embryo ID is shown at the bottom of each picture. The three embryos # cb09 04, cb17 05 and cb15 01 are those also displayed in Figure 4D. There are reproduced here for easier comparison. The two outlier embryos showing staining in distal limb cells are shown with a yellow asterisk.

**Supplementary Figure 4-3.**
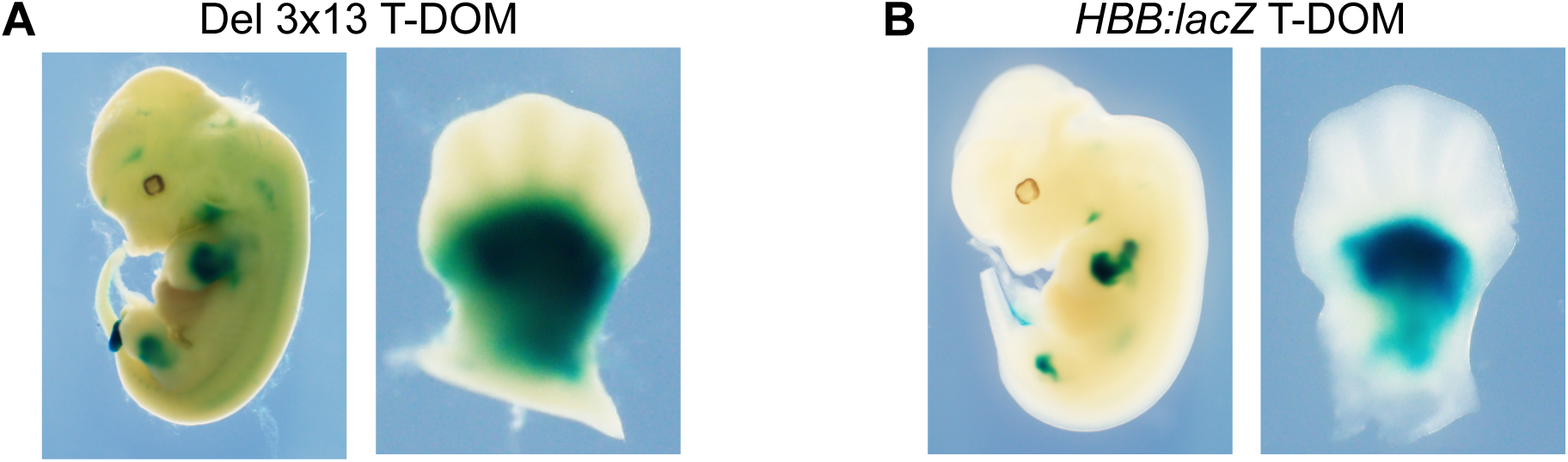
*LacZ* staining pattern in control transgenes inserted into T-DOM. **A**. The *LacZ* staining in a control embryo containing the Del 3x13 variant of the II1 enhancer. This variant contains the same targeting construct as the II1 T-DOM 542 allele, but the II1 enhancer has been replaced with the same variant used to produce the randomly integrated Del 3x13 (Figure 4D). The staining is very strong in the proximal limb and completely absent in the distal limb. This corroborates the observation from Figure 4, that the three HOX13 binding sites in II1 are necessary for the distal limb staining either as a randomly integrated transgene or a targeted insertion transgene in the T-DOM. **B**. The *LacZ* staining in a control embryo that does not contain the II1 enhancer element. This transgene contains the same targeting construct as the II1 T-DOM 542, but without the enhancer. The staining is very strong in the proximal limb and completely absent in the distal limb.

**Supplementary Figure 5.**
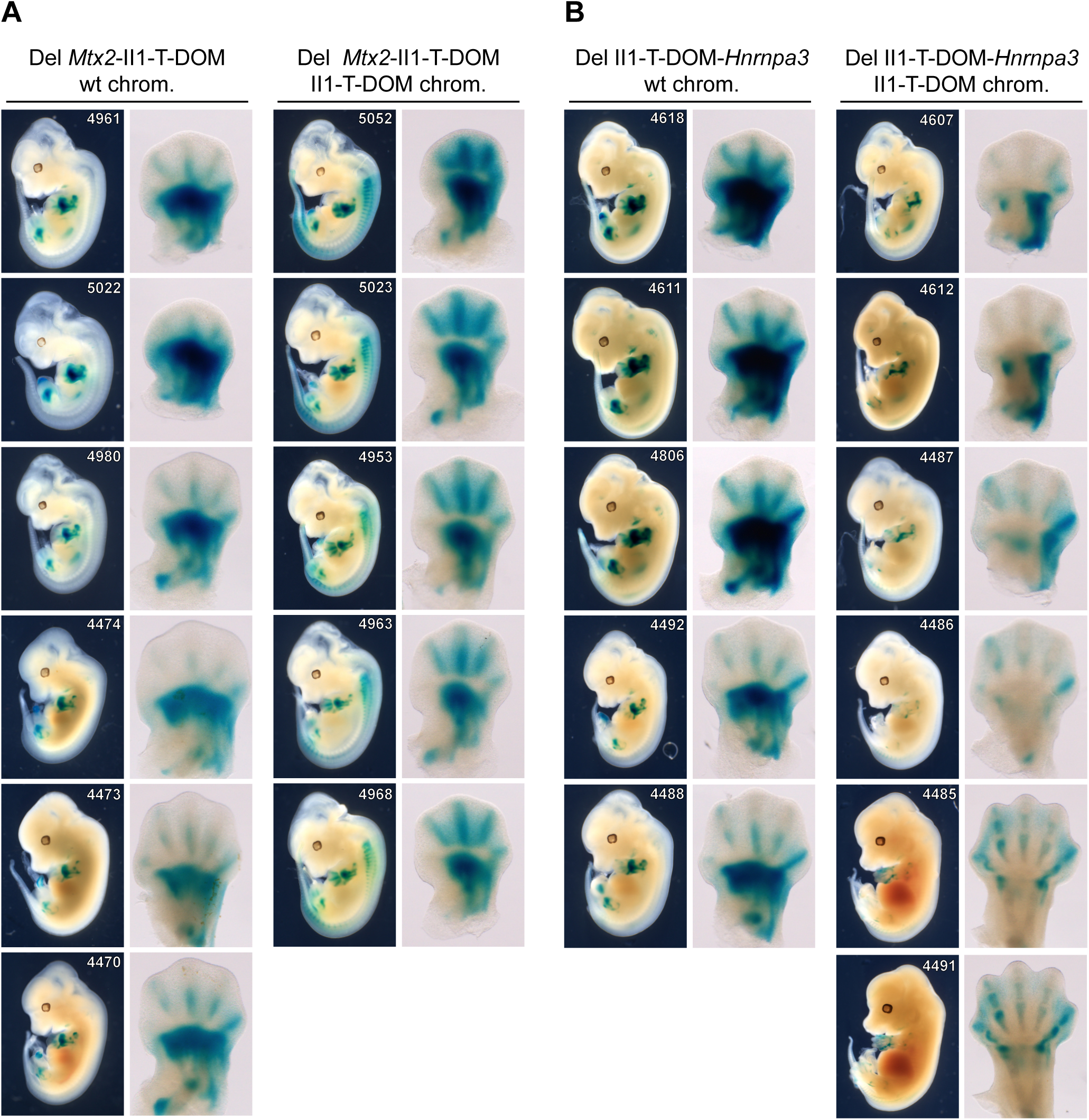
Photos of all embryos carrying large deletions flanking the insertion of the of II1 enhancer-reporter construct within T-DOM (see Figure 5B, C). All embryos were genotyped for the expected deletion. All embryos with the expected deletions are represented here unless they produce ambiguous PCR results or were mosaic for the deletion. The embryos #4470, 5023, 4492, 4491 from Figure 5 are reproduced here for easier comparison.

**Supplementary Figure 6.**
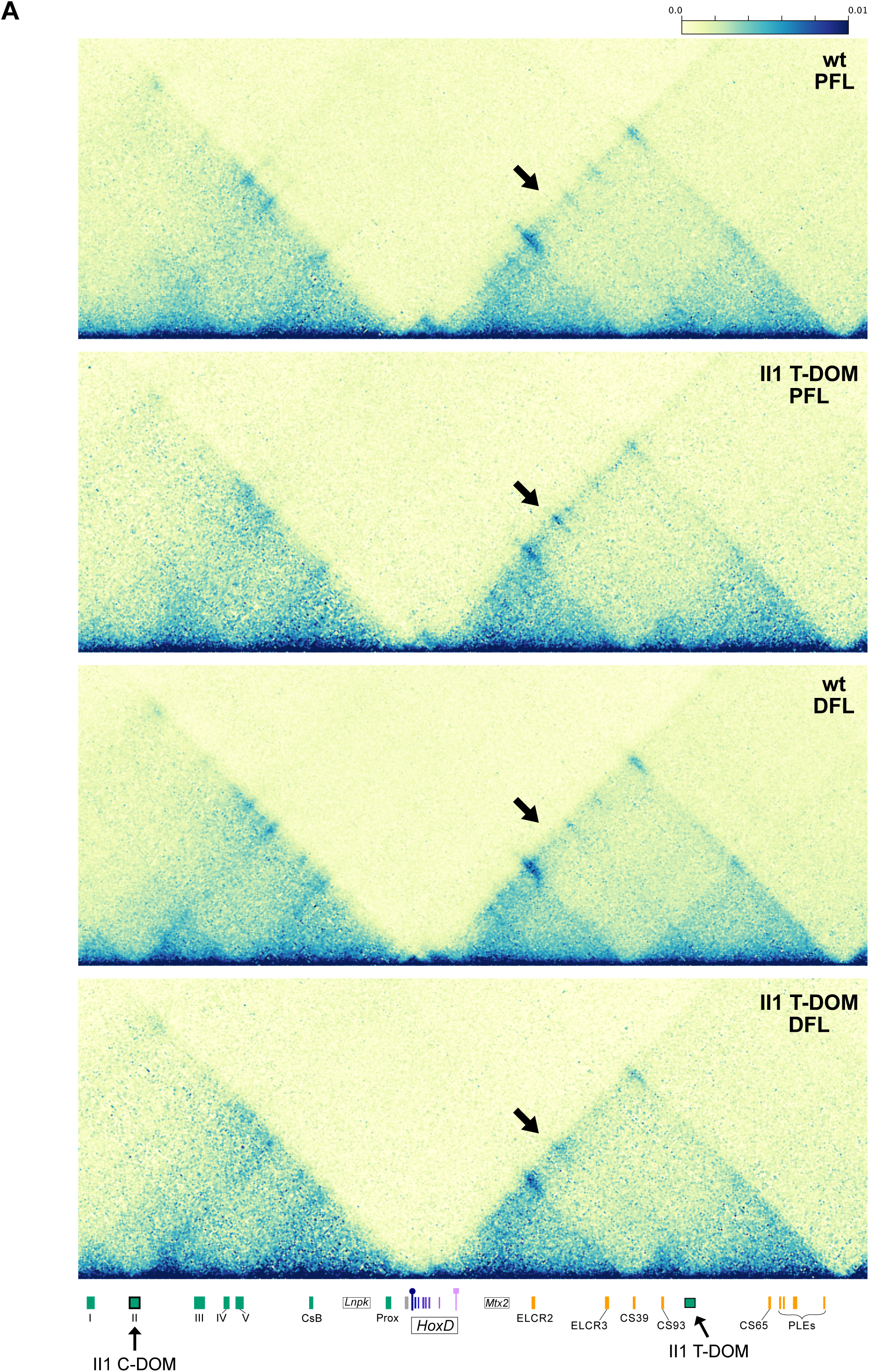
Capture Hi-C maps. Capture Hi-C maps covering the entire *HoxD* locus for wild type (wt) and the II1 enhancer recombined within T-DOM (II1 T-DOM), using both proximal (PFL) and distal (DFL) forelimb cells samples (mm10 chr2:73950000-75655000). The new contacts formed between the II1 enhancer-promoter sequence and the *Hoxd* gene cluster are indicated by the black arrow. The contacts were scored in both PFL and DFL cells, yet with a difference in resolution, being more diffuse in the DFL cells (see Figure 6). The black arrows below points to the location of the II1 enhancer within C-DOM and the transgene integration site within T-DOM. The *HoxD* cluster position is indicated at the center. The *Hoxd13* gene is indicated by a purple pin with a circle and the *Hoxd1* is indicated with a square.

